# A global analysis of climate-driven reversal risks in forests

**DOI:** 10.64898/2026.06.19.733404

**Authors:** Chao Wu, Michael L. Goulden, James T. Randerson, Anna T. Trugman, Jonathan A. Wang, Linqing Yang, Nezha Acil, Susan C. Cook-Patton, Danny Cullenward, Steven J. Davis, Christopher A. Williams, William R. L. Anderegg

## Abstract

The integrity of forest-based climate solutions and carbon credits requires persistent carbon storage, but climate change is increasing the risk of natural disturbances that release carbon back into the atmosphere. Using global satellite data, disturbance modeling, and machine learning, we provide the first spatially explicit and scenario-based maps of long-term probability of carbon loss in global forests under different disturbance severities and climate scenarios. We find that North American conifer forests, tropical rainforests, and Asian (sub)tropical dry forests face the greatest risks, and that Eurasian temperate forests, African (sub)tropical dry forests face the lowest. Globally, the likelihood of reversals over 100 years is 31%−42% across all scenarios. Our work helps to maximize the benefits of forest-based climate solutions by informing more strategic project placement and more robust reversal-risk compensation mechanisms, such as buffer pools, and highlights critical additional science to better understand and manage risks of these essential climate solutions.

**Plain Language Summary:** Forests can help slow and lessen climate impacts. However, in places this benefit is becoming less reliable as climate change increases natural disturbances such as wildfires, drought, storms, and insect outbreaks, which can release stored carbon back into the atmosphere. In this study, we created the first scenario-based global maps of risks and found that the risk of carbon loss is widespread and highly variable across regions, with especially high vulnerability in North American conifer forests, tropical rainforests, and Asian tropical and subtropical dry forests. Our study highlights the importance of considering disturbance risks when siting forest projects for climate mitigation, and developing protocols for carbon markets, such as in voluntary programs and under the UNFCCC Paris Agreement.

**Key Points:**

- A demographic model framework estimates the reversal risk from natural disturbances over 100 years in global forests
- Spatially explicit maps under different severity scenarios show variation in the integrated 100-year risk of carbon reversal
- Spatially explicit maps estimate the required buffer pool needed to compensate for disturbance-driven reversals in global forests

## 1 Introduction

Forests play a major role in global carbon cycling, acting as a substantial carbon sink [Bonan, 2008; Pan et al., 2024]. Forests could further contribute to climate mitigation in the 21^st^ century through human interventions to increase forest carbon storage via ‘nature-based climate solutions’ (NbCS) [Anderegg et al., 2020; Griscom et al., 2017; Roe et al., 2019; Seddon, 2022; Seidl et al., 2017]. Prominent forest NbCS include avoided deforestation, reforestation/afforestation, improved forest management, and agroforestry [Buma et al., 2024; Ellis et al., 2024; Griscom et al., 2017]. However, climate-sensitive disturbances, such as wildfire, drought, and pests, increasingly threaten forest carbon stocks [Anderegg et al., 2020; Brodribb et al., 2020; Byrne et al., 2024; Seidl et al., 2017; Wu et al., 2023; Wu et al., 2026] and are already having major impacts on forest carbon in many regions, potentially associated with a weakening terrestrial carbon sink [Byrne et al., 2024; Hudiburg et al., 2023; Randerson et al., 2025].

A wide range of policy approaches can support and catalyze forest NbCS, but to deliver climate mitigation benefits, they must be scientifically robust in a number of key criteria [Anderegg et al., 2025b; Ellis et al., 2024; Haya et al., 2023]. The durability of the carbon stored is a central criteria, as storage needs to be maintained almost a century or longer to deliver effective climate mitigation impact [Anderegg et al., 2025b]. Carbon credit markets are a prominent way of funding forest NbCS and primarily strive to achieve durability using a risk-transfer (e.g., insurance-like) mechanism called ‘buffer pools’, where a fraction of issued carbon credits is held in reserve to compensate for carbon losses from natural or other unavoidable causes over a defined period of time [Badgley et al., 2022; Galik and Jackson, 2009]. Yet, most current buffer pools do not vary through space, consider climate change acceleration of risks, and/or come from independent, peer-reviewed estimates [Anderegg et al., 2025a; Anderegg et al., 2025b; Badgley et al., 2022; Haya et al., 2023; Hurteau et al., 2013; Wu et al., 2023]. Without this information there is a risk that projects are inappropriately sited in places that are too risky, and that buffer pools will be too small. Thus, we urgently need data-constrained estimates of the magnitude and spatial patterns of potential climate-driven carbon losses (frequently termed ‘reversals’) and the size of buffer pools needed to effectively maintain storage.

Mapping the magnitude and spatial pattern of risk requires integrating several related elements: (1) the amount of exposed carbon, (2) the probability that exposed carbon will be disturbed, (3) the likely magnitude of carbon lost if disturbed, and (4) the amount of carbon recovery after disturbance within a given time window (e.g., monitoring or crediting period). Not only do these factors vary through space, they are also shifting as the climate changes. Furthermore, the reversal risk analysis needs to align with the requirements of the applicable carbon crediting mechanism, is 100 years in the Verra Verified Carbon Standard program and California compliance market [Anderegg et al., 2025a; Haya et al., 2023; Michaelowa et al., 2025; Wu et al., 2026], though we note that longer commitments of up to 1000 years or more may be necessary to compensate for the warming effects of carbon dioxide emissions [Anderegg et al., 2025b; Brunner et al., 2024]

To address each of these factors, we combined remotely sensed forest biomass and stand age data, and historical and future climate and disturbance data, with machine learning models and a demographic modeling approach that directly modeled forest carbon stocks per decade, accounting for both disturbance and regrowth. The result is the first spatially explicit and scenario-based estimates of the probability of at least one reversal event over 100 years (i.e., “integrated 100-year probability”) due to natural disturbances for global forests. We show results for different disturbance severity and climate scenarios (see a flowchart figure in Fig. S1). To account for uncertainty in forest carbon stock data, we determined a “reversal” as a 10% or more decrease in net live biomass between each future decade relative to the historical but also explore 0% as a threshold as a sensitivity analysis, Fig. S2. We next provided spatially explicit map of the buffer pool needed to cover carbon lost to natural disturbances, assuming all lost carbon would be covered by the buffer pool (although see Text S1 for an alternate approach). To account for future climate conditions, we use Shared Socioeconomic Pathways (SSP) 2-4.5 data, which represents a moderate-emissions scenario with moderate climate change that is most consistent with current climate policy commitments [Climate Action Tracker, 2023; O’Neill et al., 2016]; we also included a high-emissions scenario SSP5-8.5 as a supplement. The end result is a series of scenario-based maps that can help inform more stable locations for investment and more robust compensation mechanisms, and help advance higher-integrity carbon markets and climate policy frameworks.

## 2 Materials and Methods

### 2.1 Global-scale demographic models to calculate carbon reversal probability

We built a global-scale demographic ‘growth-mortality’ model that directly estimates forest aboveground biomass (AGB) changes in response to disturbance and subsequent regrowth, building on our previous regional-scale effort [Wu et al., 2023]. There were four steps to this analysis that involved (1) developing forest growth curves, (2) predicting the probability of disturbance over 10-year intervals, (3) estimating the amount of carbon lost from disturbance, and (4) calculating the integrated 100-year probability of carbon reversal. Each step is described below.

The first step involved developing forest growth curves per 8km grid, running the model separately for each of 19 broad forest ecoregions (e.g., moist tropical forest in America; [Xu et al., 2021]). The curves fit AGB data (from ESA CCI AGB V4.0, [Santoro and Cartus, 2023]) as a function of stand age (from [Santoro and Cartus, 2023]), as well as mean annual temperature (MAT) and precipitation (MAP) (ERA5; [Hersbach et al., 2018]). To determine the best curve, we tested 10 different growth functions with gamma-distributed noise, optimized the parameters using maximum likelihood as defined by the gamma distribution, and selected the best-fitting model (Table S1). If no curve achieved a R^2^ ≥ 0.15 at the ecoregional level (an acceptable fitting performance based on similar studies in the literature [Anderegg et al., 2022b; Wu et al., 2023; Zhu et al., 2018]), then we reran the model at the biome level (from [Olson et al., 2001]) and used the best biome-level curve for each 8-km grid (see Table S2 for fitting performance). We used 70% of the total grid cells to fit the model, with the remaining 30% kept for testing.

The second step involved modeling the probability of stand-replacing disturbances. We first mapped stand-replacing disturbances using Global Forest Watch data [Hansen et al., 2013] for 2001-2022 at an original half-degree resolution and further regridded to an 8 km × 8 km grid. To focus on natural disturbances, we removed tree cover loss resulting from human land use change (similar to [Anderegg et al., 2022a]) and heavy forest management (from [Lesiv et al., 2022], see Fig. S3). Thus the remaining areas are assumed to represent a mix of natural disturbances (e.g., wildfires, insect and disease outbreak, and windthrow). We converted these to annual disturbance probability (%/yr; by dividing by grid area of 8km-resolution and period length (i.e., 22 years)). We then built a random forest model that linked annual disturbance probability based on historical mean annual temperature, mean annual precipitation (ERA5; [Hersbach et al., 2018]), AGB [Santoro and Cartus, 2023], and biome [Olson et al., 2001], which explained 43.3% of variance. To then examine disturbance probabilities under future climate, we reran the random forest model with projected temperature and precipitation values per decade between 2010 and 2100. We used a ‘pseudo-global warming’ approach to calculate the projected changes in future climate while avoiding any potential absolute bias in Earth System Models (ESMs) over the historical periods, because it applies the delta between future and historical from the ESMs (in the CMIP6 in this analysis) to the observed historical meteorological data (ERA5). These projected values represent a multi-model mean across 22 ESMs that represent the spread of climate projections across or within model families, running a middle-of-the-road and a high-emissions future climate scenario (SSP2-4.5 and SSP5-8.5) [Anderegg et al., 2022a] (Table S3). Finally, we scaled disturbance probabilities from an annual probability to a decadal probability, using a binomial distribution algorithm. We then spatially interpolated the predicted disturbance probability back into areas previously excluded due to heavy anthropogenic disturbance using inverse distance weighted interpolation to provide wall-to-wall reversal probability maps. Several studies have identified inconsistencies in the forest change time series from GFC data, especially starting from year 2015, partly due to the update of the GFC algorithm [Palahí et al., 2021]. Therefore, we conducted a sensitivity analysis on the disturbance detection periods and found that using either 2002–2014 (used in Anderegg et al. [2022a]) or 2001–2022 (this study), yielded broadly similar magnitudes and spatial patterns in both historical and predicted disturbance probability (Text S2 and Fig. S4).

The third step involved determining how much carbon is lost upon disturbance. Although the map of disturbance probability was based on stand-clearing disturbances, many disturbances are more partial or diffuse and do not release all the carbon stock. The amount of carbon lost to these partial disturbances shows large variation across biomes and severity gradient [Pugh et al., 2019; Thom and Seidl, 2016]. This makes our disturbance probability map conservative (i.e., an under-estimate) of the overall likelihood of disturbance. To capture variation in carbon loss we relied on existing literature syntheses. For boreal and temperate forests, stand-replacing disturbances (wildfires, wind, and bark beetles) caused a decrease in ecosystem total carbon by 38.5% and aboveground live carbon by 91.3% on average [Thom and Seidl, 2016]. We treated 38.5% and 91.3% as lower and upper bounds for AGB loss. Moderate severity wildfires tended to cause a decrease in basal area (a linearly-related proxy for aboveground live carbon) by 71.6% [Whittier and Gray, 2016], so we designated 71.6% AGB loss as a moderate scenario. For tropical forests, windthrow and wildfires have been shown to cause a percent of biomass loss of about 45% and 55% on average, with a maximum loss of about 65% from wildfires [Bullock and Woodcock, 2021], which we designated as the low, moderate, and high AGB loss scenarios, respectively. However, we acknowledge that these severity values were based on initial literature assessments. We further assumed that the stand age (but not biomass) would change back to zero after experiencing disturbances following previous literature [Pugh et al., 2019], though we acknowledge that disturbance severity will determine if the stand age functionally ‘resets’ back to zero. We primarily visualize three illustrative severity scenarios, namely ‘low’ severity, ‘moderate’ severity, and ‘high’ severity for all biomes under SSP2-4.5 (Table S4). However, we include the scenario-based results from full combinations of severity of AGB loss across biomass (i.e., nine scenarios) and the other climate scenario in Table S5.

Finally, we combined each of these three steps above, resampling each to a common 8 km × 8 km grid and built a demographic growth-mortality model at this resolution to calculate the integrated 100-year probability of carbon reversal due to natural disturbances. We ran the model only for areas that had >10% tree cover in the year 2000 [Anderegg et al., 2022a]. The reversal simulations were structured as follows. We first scaled forest age in all grids to the year 2010 and then predicted AGB in subsequent decades (that is, the 2010s, 2020s,…, 2090s) for each of the 19 ecoregions independently using the best fitted growth curve. To capture the stochasticity inherent in disturbance probabilities, we conducted a Monte Carlo simulation that generated 100 randomly distributed binomial disturbance occurrences (0 or 1) based on decadal disturbance probabilities for each grid cell. For each run, if there was a disturbance, we reduced AGB by the relevant severity percentage and reset the regrowth back to age zero. We then calculated the integrated 100-year probability of carbon reversal caused by natural disturbances for each 8-km grid following equations 1 and 2.

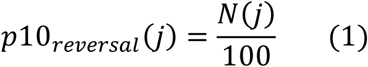

Where *p*10_*reversal*_ (*j*) is the decadal carbon reversal probability due to natural disturbances for a given 8 km-grid, *j* represents the decade (2020s, 2030s, …, 2090s), *N*(*j*) is the number of simulations from the total of 100 where (1) a disturbance occurred and (2) the net of AGB loss and regrowth during decade *j* was 10% or lower than AGB in the 2010s. Here, we used AGB as a proxy of total live biomass carbon to identify a reversal under our 10%-threshold rule because we approximated total live biomass from AGB * (1 + root:shoot). The integrated 100-year carbon reversal probability (*p*100_*reversal*_, %) due to natural disturbances was calculated using equation (2).

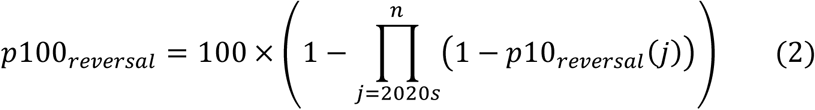

Where *n* represents 10 decades (eight decades from the 2020s to 2090s and then a repeat of 2080 and 2090, as there are no climate data for 2100 and 2110) to integrate probabilities over a full 100 years.

### 2.2 Maps of the buffer pools to compensate for disturbance-driven reversals in global forests

We next calculated a map of biophysical carbon loss as a key buffer pool metric for disturbance-driven reversals to directly inform policy efforts. First, we calculated wall-to-wall maps of the “expected C loss” (%) across all global forested lands at an 8-km spatial resolution which provided guidance for the percentage of live carbon predicted to be lost due to natural disturbances across a range of disturbance severities scenarios over 100 years under two future climate scenarios. We calculated these maps using the integrated 100-year carbon reversal probability, estimated above, and the disturbance severity (i.e., the ratios of carbon losses where a reversal event occurs due to a given disturbance dependent on biomes: 38.5%, 71.6%, and 91.3% for boreal and temperate forests, and 45%, 55%, and 65% for tropical forests) at an 8km grid level following equation (3).

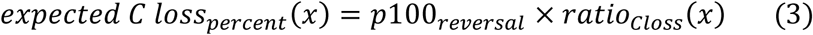

Where *ratio*_*Closs*_(*x*) represents the disturbance severity scenarios, i.e., the ratio of the carbon losses where a reversal event occurs due to a given disturbance dependent on a biome *x*.

Then, we calculated maps of the ‘required buffer pool density (tCO_2_e ha^-1^) to compensate for disturbance-driven reversals’ that provide the absolute magnitude of carbon by combining expected C loss and a map of total (aboveground and belowground) live biomass carbon exposure map (equation 4). For total live biomass carbon, we used a map of global AGB circa 2017 from the ESA CCI AGB product V4.0 at a 100 m spatial resolution [Santoro and Cartus, 2023] and a spatially explicit estimate of root:shoot ratio dataset at ∼1km spatial resolution [Huang et al., 2021]. Spatially, we resampled all datasets to a common 8 km × 8 km grid by a bilinear interpolation method.

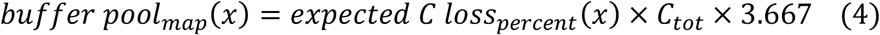

Where *buffer pool*_*map*_(*x*) is the maps of the required buffer pool density (tCO_2_e ha^-1^) to compensate for disturbance-driven reversals across global forests, *expected C loss*_*percent*_ (*x*) is the same as in equation 3, *C*_*tot*_ is the total (aboveground and belowground) live biomass carbon stocks (t ha^-1^), and 3.667 is a scaling factor from carbon to CO_2_e. We note that standing dead and soil carbon are not directly included in this analysis (see Text S3). However, future studies may consider these pools in forest carbon protocols to align with the critical role of these pools in global carbon cycling.

We note that translating tonne-based results from equation 4 into a parameter expressed in terms of the percentage of credit carbon stocks, as is commonly used in the design of buffer pools in real-world carbon crediting programs, depends on the technical details of carbon crediting methodologies and individual projects. We discuss different approaches taken in real-world carbon markets and illustrate how to make these additional calculations under one of two common accounting paradigms in use today (see Text S1).

## 3 Results

### 3.1. Carbon reversal probabilities from natural disturbances in global forests

When we mapped the integrated 100-year probability of carbon reversal due to natural disturbances at 8-km resolution, we found substantial variation in the likelihood of a reversal through space. We evaluated nine different scenarios that differed in how severe disturbances were and how those severities were distributed under each of two future climate scenarios. We visualized three primary severity levels (globally low, moderate, or high severity; Fig. 1) under SSP2-4.5 in the main text but present full scenario-based estimates in Figs. S5 and S6. In general, the 100-year probability of reversal increased with disturbance severity, and dry forests in the Amazon, North America, southern China, and Congo rainforests were predicted to be exposed to higher reversal probabilities, especially in higher severity scenarios, and the lowest risk in Eurasian temperate forests, African tropical and subtropical dry forests. Overall, the average global 100-year reversal probability was 31%−42% (depending on the severity and future climate scenarios).

**Figure 1.**
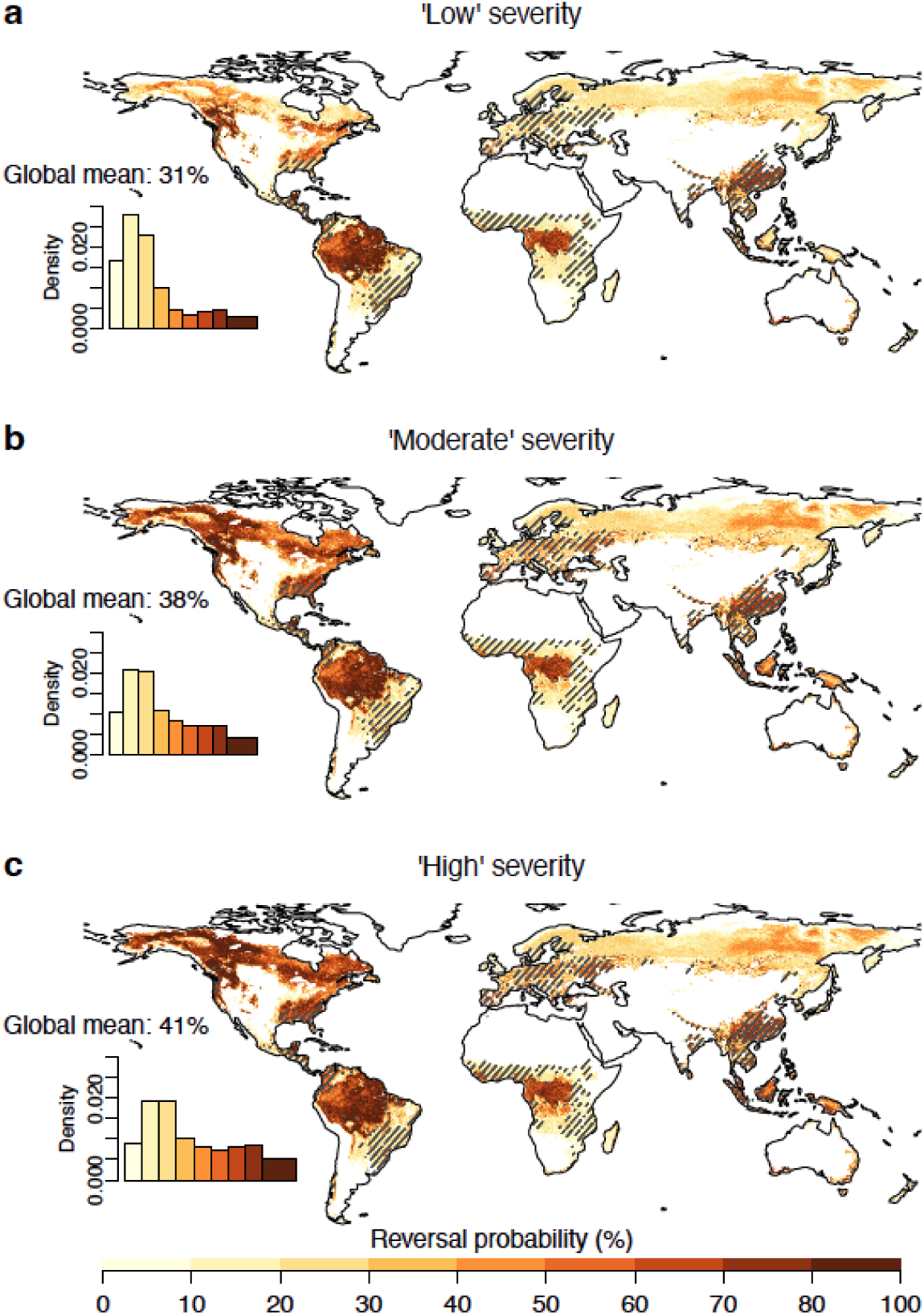
Spatial patterns of the integrated 100-year carbon reversal probability in global forests due to disturbances under low (a), moderate (b), and high (c) severity scenarios in SSP2-4.5. Global weighted mean reversal probability by grid area is indicated. Hatched areas indicate the areas with heavily managed forests where probability estimates were interpolated. The inset figures show the kernel density estimates across all forested 8-km grid cells.

Compared to tropical forests, boreal and temperate forests were a larger contributor to uncertainty in reversal risk estimates due to poorly constrained estimates of severity of carbon loss in those forests. In addition, reversal likelihood exhibited very similar spatial patterns under the two climate scenarios (Figs. S5 and S6), suggesting that uncertainty associated with disturbance severity was substantially greater than that associated with climate change.

### 3.2 Divergent reversal probabilities across different forest ecoregions

We also examined 100-year probabilities of reversal by ecoregion (Figs. 2 and S7). Of the 19 ecoregions, 5 had larger reversal probabilities than the global mean under the low severity scenario, 9 under the moderate severity scenario, and 10 in the high severity scenarios. These patterns were broadly consistent across climate scenarios. Notably, the divergence in reversal probabilities across ecoregions (represented as a ratio of maximum and minimum risks) in lower severity scenarios (73% for tropical moist forests in the Americas vs. 12% for North American Tundra) was larger than that in higher severity scenarios (79% for North American conifer forests vs. 15% for Tropical and subtropical other vegetation in Africa). These indicate that reversal risks are more localized under the low severity scenario, but become increasingly predominant under higher severity scenarios.

**Figure 2.**
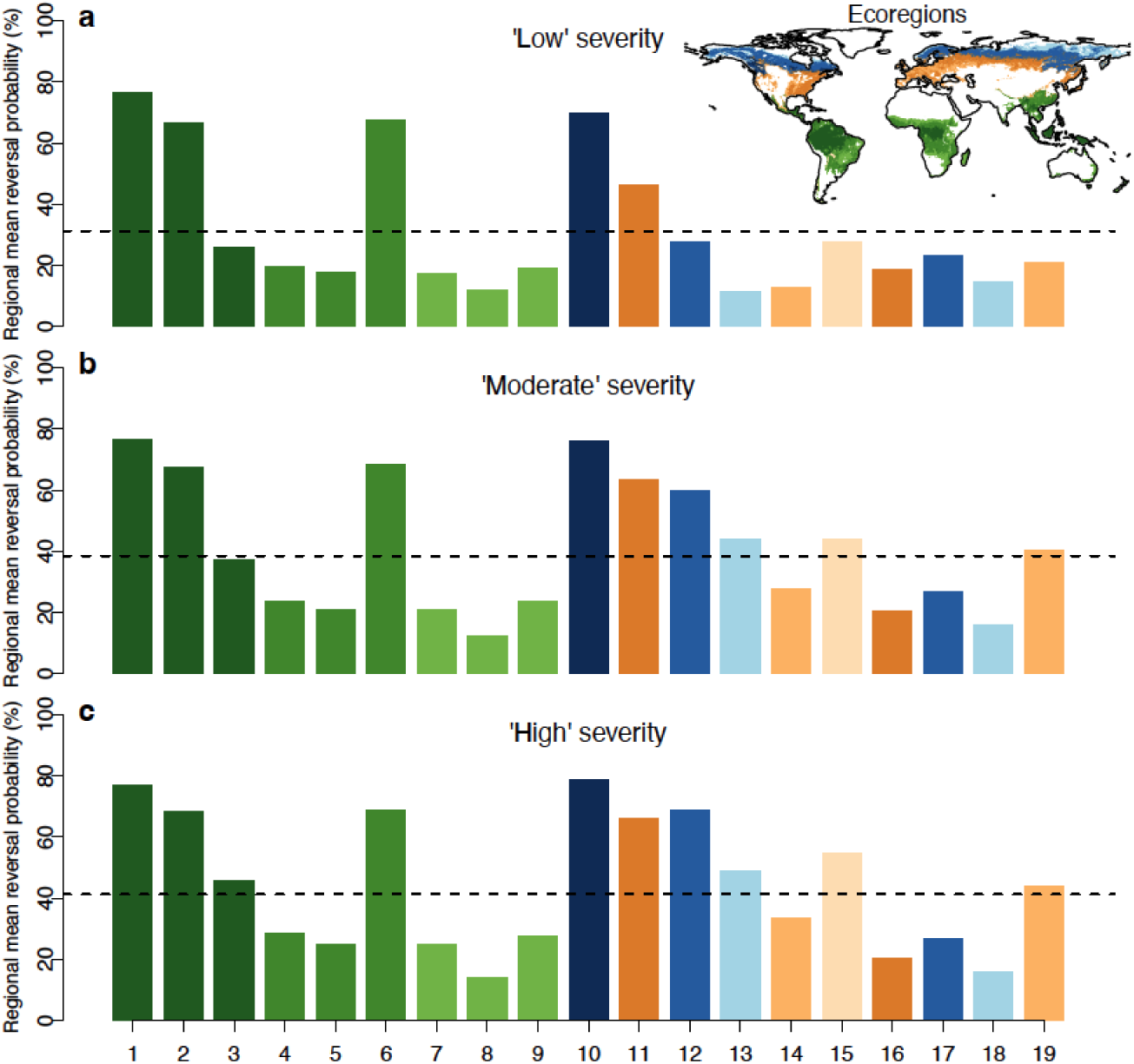
Regional mean probabilities of carbon reversal are provided at a forest ecoregion level under three major severity levels: low (a), moderate (b), and high (c) in SSP2-4.5. The 19-ecoregion used in this study is based on Xu et al. [2021]. The labels from 1 to 19 represnet the ecoregions of 1: Moist Tropical Forest (Americas), 2: Moist Tropical Forest (Africa), 3: Moist Tropical Forest (Asia, 4: Tropical & Subtropical Dry Forest, Shrubland (Americas), 5: Tropical & Subtropical Dry Forest, Shrubland (Africa), 6: Tropical & Subtropical Dry Forest, Shrubland (Asia), 7: Tropical & Subtropical Other Vegetation (Americas), 8: Tropical & Subtropical Other Vegetation (Africa), 9: Tropical & Subtropical Other Vegetation (Asia), 10: Conifer Forest (North America), 11: Temperate Forest, Shrubland (North America), 12: Boreal Forest, Shrubland (North America), 13: Tundra (North America), 14: Temperate Other Vegetation (North America), 15: Southern Forest (South America), 16: Temperate Forest, Shrubland (Eurasia), 17: Boreal Forest, Shrubland (Eurasia), 18: Tundra (Eurasia), 19: Temperate Other Vegetation (Eurasia). The horizontal line is the global mean reversal probability in Fig. 1.

Across ecoregions, North American conifer forests, tropical moist forests in the Americas and Africa, Asian tropical and subtropical dry forests, North American boreal, and temperate forests were at high probability of climate-driven reversals, whereas Eurasian temperate forests, African tropical and subtropical dry forests, and other non-forests-dominated ecoregions were at generally lower reversal probability. Interestingly, moist tropical forest risk varied by continent and was higher in the Americas and Africa, but lower in Asia.

### 3.3 Buffer pool needed to compensate for disturbance-driven reversals in global forests

Finally, we calculated the global maps of the ‘required buffer pool density (tCO_2_e ha^-1^) to compensate for disturbance-driven reversals’ (Figs. 3, S8, S9; for an alternate formulation, see Figs. S10 and S11 and Text S1) to improve forest carbon offset programs and to inform project selection to better cope with escalating climate-driven durability risks. Our results highlighted that areas with high carbon density (e.g., tropical rainforests) need higher buffer pool densities given the high amounts of carbon potentially exposed to disturbance. We also found that North American conifer forests and Eurasian boreal forests needed higher buffer pool densities given the greater probability of a reversal in those locations. However, we also found that disturbance severity, which is currently poorly constrained in the literature, has a major impact on the required buffer pool density, highlighting a key area of needed additional research.

**Figure 3.**
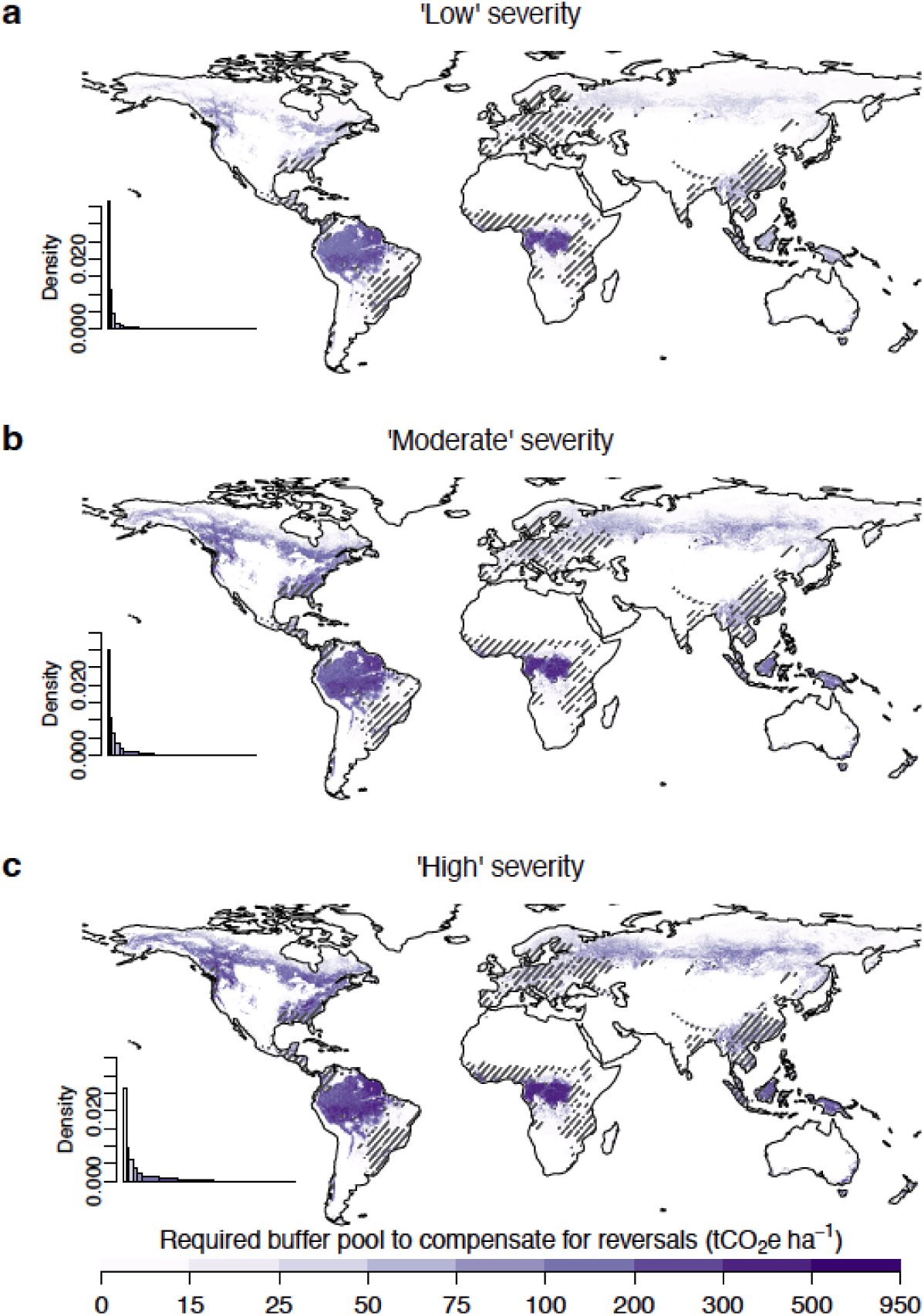
Spatial patterns of the required buffer pool density (tCO_2_e ha^-1^) to compensate for disturbance-driven reversals under low (a), moderate (b), and high (c) severity scenarios in SSP2-4.5. Hatched areas indicate the areas with heavily modified forests. The inset figures show the kernel density estimates across all forested 8-km grid cells.

## 4 Discusion

NbCS projects can only act as climate solutions if they are sufficiently durable, and yet escalating climate change threatens that durability. Buffer pools can help ensure durability, but only if they are sufficiently sized. Here, we provided the first global spatially explicit and scenario-based maps, at 8-km resolution, of the 100-year probability of carbon reversal and associated buffer pool contributions to cover carbon loss (expressed in tCO_2_e ha^-1^). The scenarios help to capture the range of potential outcomes, and the maps can help prioritize projects in places that are less likely to be disturbed and inform the design and update of NbCS crediting programs. Our analysis indicates that North American conifer forests and tropical rainforests are at high likelihood of reversals, consistent with recent observations of widespread carbon losses in western and southern North America due to wildfires and severe drought [Kannenberg et al., 2021; Michaelian et al., 2011; Schwalm et al., 2012; Wang et al., 2022; Wu et al., 2026] and in tropical rainforests driven by drought-driven tree mortality and climate variability [Forzieri et al., 2022; Gatti et al., 2021; Hubau et al., 2020; Nolte et al., 2023; Tavares et al., 2023]. These ecosystems combine large carbon stocks with high exposure to stand-replacing disturbances, increasing the likelihood of substantial and persistent carbon losses [Pugh et al., 2019; Seidl et al., 2016]. In contrast, Eurasian temperate forests and African tropical and subtropical dry forests exhibit lower reversal probabilities, likely reflecting lower disturbance impacts and/or smaller carbon losses following disturbance. Although reversal probabilities in Eurasian boreal forests were estimated to be relatively low under baseline assumptions, their large carbon stocks make them particularly sensitive to disturbance severity. As a result, high-severity standing-replacing disturbances [Anderegg et al., 2022a; Seidl et al., 2020] can release substantial amounts of carbon and markedly increase reversal risk, making these forests a potential hotspot for larger buffer pool contributions under high-severity scenarios.

Our study provides usable and urgently needed data for active and ongoing policy discussions, but uncertainty is inherent in any future projections. Thus, our study also highlights key uncertainties, limitations, next steps, and research opportunities. Uncertainty arises from the quality of biomass and stand-age data, future climate and disturbance projections, and particularly from coarsely constrained estimates of disturbance severity (Table S6). In this analysis, we used a demographic modeling approach to model the dynamics of forest carbon, including critical tree growth, regrowth, and mortality under climate change, although the overall fitting performance varies across ecoregions (Table S2). Future work could leverage mechanistic land surface models, which include additional plant physiological and ecological processes but have the trade-off of being more challenging to parameterize and run at a finer spatial resolution. Further improvements should refine estimates of the amount of exposed carbon (exposure), the risk that exposed carbon will be disturbed (hazard), the likely magnitude of carbon lost if disturbed (vulnerability), and the amount of carbon recovery after disturbance. For example, here we combined all stand-clearing disturbance types in our analyses and used literature-based disturbance severity numbers for major biomes to construct our severity scenarios. However, disturbance severity remains poorly constrained by data in many regions worldwide. We found that variation in reversal-risk estimates across the nine severity scenarios was substantially larger than that across future climate scenarios. This suggests that disturbance responses to climate change are relatively weak in our disturbance model. An improved representation of different disturbance agents (and their sensitivity to climate change), as well as inclusion of more mechanistic vegetation dynamics (e.g., the degree of successful regeneration and regrowth post-disturbance), and higher resolution (downscaled) climate projection are greatly needed. Given these future research needs, our work is best presented as an integrated framework for mapping risks, that provides a best-available estimate based on illustrative and reasonable scenarios. This provides an important initial step to inform policy, and future work can improve model representation and observational constraints to better quantify global reversal risk.

Our global reversal probability and buffer pool maps have the potential to substantially improve tools designed to estimate risks. Currently, the vast majority of buffer pool protocols in voluntary and compliance carbon markets are not based on independent, rigorous estimates of reversal risk [Anderegg et al., 2025b; Badgley, 2024; Badgley et al., 2022; Sanders-DeMott et al., 2025]. The high reversal probabilities and substantial buffer pool densities we observed in some locations, suggest that buffer pool contributions will need to increase especially in higher risk and higher carbon areas. Currently, buffer contributions are quite low. For example, an analysis of 67 REDD+ projects under Verra’s previous protocols contributed a mean of 2% of credits to buffer pools to insure against all natural risks [Haya et al., 2023] and ∼12% overall for all social and natural risks. When we follow the same accounting paradigm used in Verra’s program, our results suggest the need for a 24% buffer pool contribution on average for all natural risks combined (under the ‘moderate severity’ scenario Fig. S10). Crucially, evidence-based maps of buffer pool densities that capture the substantial spatial variation in risks provide a strong incentive for more strategic carbon project placement to reduce risks. To facilitate and accelerate use and uptake of these solutions-focused buffer pool metrics, we have also built a web tool for display and downloading of our key policy-relevant datasets [a web URL will be provided when the paper is accepted]. However, the precise method for buffer pool construction varies across standards and protocols which will require some customization of the results we present here (see Text S1 for further discussion).

## 5 Conclusions

We developed a demographic model that can model forest growth, mortality, and disturbance dynamics, and applied it to estimate the 100-year probability of carbon reversal from natural disturbances across global forests under a changing climate. Our results provide estimates of buffer pool contributions required for forest carbon offset projects to maintain the durability of NbCS, while also highlighting substantial uncertainties that remain in current assessments and key steps needed for future research. We provided spatially explicit and scenario-based maps of long-term carbon reversal probabilities and buffer pool densities that could be useful for informing the evidence-based development or revision of buffer pools in real-world carbon crediting programs. Recent and near future expected advances in high-resolution Earth forest monitoring, large-scale inventory networks, and mechanistic disturbance modeling will further help constrain the current uncertainties and overcome challenges in quantifying climate risks and buffer pools.

## Acknowledgments

C.W. acknowledges support from the Tsinghua University Dushi Program, the David and Lucille Packard Foundation, and the Wilkes Center for Climate Science and Policy at the University of Utah. W.R.L.A. acknowledges support from the David and Lucille Packard Foundation and US National Science Foundation grants 2003017, 2044937, 2330582, and 2519523 as well as the Alan T. Waterman award IOS-2325700. D.C. is a member of the Paris Agreement Crediting Mechanism’s Methodological Expert Panel and California’s carbon market advisory committee, but does not speak for either group here. JTR received funding support from the US. Dept of Energy Office of Science RUBISCO Science Focus Area and NASA’s Carbon Monitoring System program. The Bezos Earth Fund supported S.C.P’s time on this work. A.T.T acknowledges support from the US NSF Grant 2003205, the Gordon and Betty Moore Foundation Grant 11974 and 13283, the USDI Park Service Award No. P24AC00910 and P24AC01425, and CALFIRE Forest Health Research Program Grant 60164685. N.A. was financially supported by the Natural Environment Research Council (NERC, NE/Y006216/1).

## Open Research

All the data used for creating the main figures in this study will be published when the paper is accepted. CMIP6 data were downloaded from https://esgf-node.llnl.gov/search/cmip6/. The AGB data was from the ESA CCI AGB product V4.0 (https://dx.doi.org/10.5285/95913ffb6467447ca72c4e9d8cf30501). The 19-ecoregion division (broad forest groups/types) was accessed from https://doi.org/10.5281/zenodo.4161694. The 13-biome division was accessed from https://doi.org/10.1641/0006-3568(2001)051[0933:TEOTWA]2.0.CO;2. Historical climate data was from ERA5 (https://doi.org/10.24381/cds.adbb2d47). Disturbance data were derived from the Landsat-based tree cover loss product, available at https://glad.earthengine.app/view/global-forest-change. Land cover data are available at https://zenodo.org/records/8239305. Forest management map in 2015 is available at https://zenodo.org/records/5879022.

## Supporting Information

### Text S1

Our primary analysis focused on quantifying the biophysical carbon losses due to disturbance for buffer pool construction (buffer pool density in tCO_2_e ha^-1^), which can inform buffer pool design across the different accounting paradigms present in carbon crediting mechanisms. In some mechanisms, such as California’s compliance offsetting program, any carbon lost is considered a reversal up to the point where losses exceeded credited carbon (i.e., such that reversals are limited by the subset of credited carbon storage rather than total carbon stocks). We refer to this as Accounting Paradigm 1 here. In contrast, other carbon crediting mechanisms, such as Verra’s Verified Carbon Standard (VCS), treat carbon losses differently. Under these mechanisms, a reversal is deemed to be the total carbon loss multiplied by the ratio between credited and total carbon (i.e., such that reversals are only a fraction of total carbon losses and are similarly capped at the level of total credited carbon storage). We refer to this as Accounting Paradigm 2 here.

Differences in accounting paradigms have implications for the technical design of buffer pools. Typically, forest carbon projects contribute carbon credits to a shared buffer pool, with contribution requirements expressed as a percentage of projects’ credited carbon outcomes. For some forest carbon project types, such as reforestation projects that begin with a baseline of approximately zero above-ground carbon, credited carbon may be similar or even equal to total carbon stocks. For other forest carbon project types, such as improved forest management projects that enhance carbon stocks above non-zero baseline levels, credited carbon may be only a fraction of total carbon stocks, with the fraction dependent on the details of the carbon crediting methodology and the individual project. These fractions can vary widely from [0,1].

Under Accounting Paradigm 1, translating the estimated required buffer pool density (tCO_2_e ha^-1^) to a corresponding percentage of credited carbon depends on the specific choices made in carbon crediting methodologies and by individual projects. We report our main results in terms of the estimated required buffer pool density (tCO_2_e ha^-1^) and note that the expression of these calculations in terms of a percentage of credited carbon is contingent on methodological and project-specific choices, as these choices affect the ratio of credited to total carbon stocks. Although the same choices are also relevant under Accounting Paradigm 2, they have no effect on the translation the estimated required buffer pool density (tCO_2_e ha^-1^) into a percentage of credited carbon because Accounting Paradigm 2 assigns forest carbon losses as reversals based on the same ratio of credited to total carbon stocks. Thus, under Accounting Paradigm 2, the translation of total reversals (tCO_2_e ha^-1^) as a percentage of credited carbon stocks is independent of ratio of credited to total carbon stocks. We note that the reasons to prefer one accounting approach over the other are nuanced and beyond the scope of our analysis, which seeks to inform buffer pool designs independent of their preferred accounting approach.

To illustrate how our analysis can directly inform buffer pool designs in carbon crediting mechanisms operating under Accounting Paradigm 2, we quantified the percent of credited carbon that projects would need to contribute to the buffer pool as the product of reversal probability and disturbance severity (i.e. the ‘expected C loss’) (Figs. S9 and S10). For this metric, the expected C loss was most sensitive to where reversal probabilities were predicted to be modest (e.g., Eurasian boreal forests) but could be amplified in higher severity scenarios. This suggested that, beyond climate-driven reversal probability, potential changes in disturbance severity, which are currently poorly constrained, in a changing climate have major impacts on further buffer pool percent contributions.

Similar calculations can be produced for carbon crediting mechanisms under Accounting Paradigm 1 but are contingent on how the ratio between credited carbon and total carbon stocks is determined in those mechanisms. We note that because Accounting Paradigm 1 treats a larger share of forest carbon losses as a reversal than does Accounting Paradigm 2, percentage-based buffer pool parameters will be strictly higher under Accounting Paradigm 1 relative to those calculated under Accounting Paradigm 2 (e.g., Figs. S9 and S10). Future work could identify how these methodological choices translate into percentage-based parameters for specific carbon crediting mechanisms operating under Accounting Paradigm 1.

### Text S2

Additionally, our sensitivity analysis of historical disturbance datasets indicated very similar results when comparing the GFC data for 2002-2014 versus 2001-2022, indicating our approach is likely robust to the time period of historical data used (Fig. S3). Across all 8-km grid cells, the datasets exhibited a strong agreement, with a Pearson correlation coefficient of r = 0.86 and a Spearman rank correlation of r = 0.98. When limited to forest-only grid cells, the correlations remained high (Pearson r = 0.83, Spearman r = 0.93). Ordinary least squares (OLS) regression between the two datasets yielded an R² = 0.69, a slope of 0.9, and a p-value < 0.0001, indicating a statistically significant linear relationship.

### Text S3

We note that standing dead and soil carbon are not directly included in this analysis. because i) they are relatively poorly understood and challenging to monitor change globally, ii) most carbon market protocols do not require including these pools, iii) most standing-dead trees are expected to fall down within the decadal time-step based on current literature estimates, and iv) soil carbon typically only matters in limited cases with strong soil management and is generally not considered crucial for forest NbCS. For example, a recent study estimated that soil carbon comprises only 9% of the mitigation potential in forests as a NbCS [Bossio et al., 2020].

---

**Figure S1.**
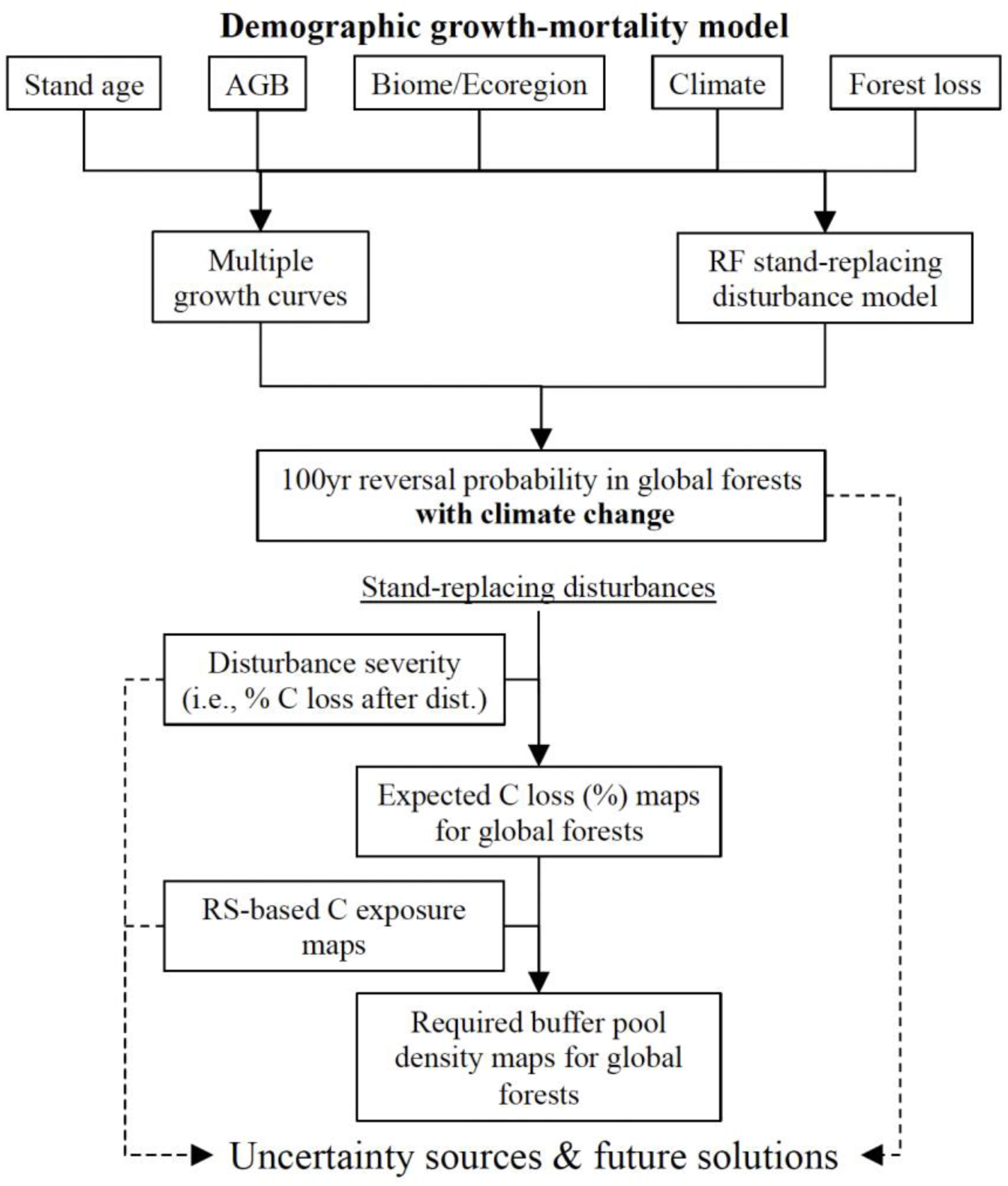
Flowchart of the overall methods used in this study.

**Figure S2.**
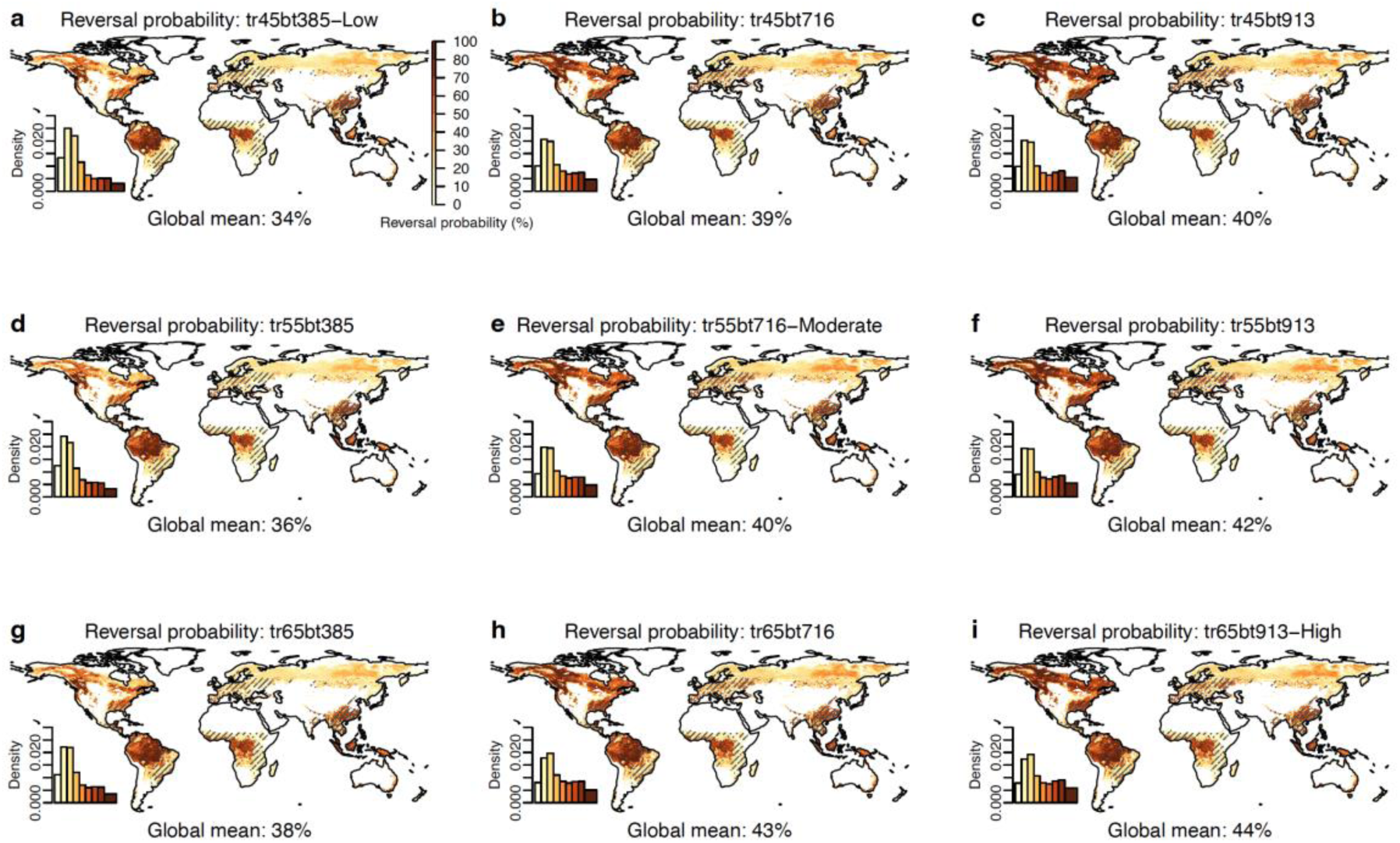
Spatial patterns of the integrated 100-year carbon reversal probability in global forests due to disturbances under the full range of 9 severity scenarios in SSP2-4.5 by combining the conditions where AGB was reduced by 38.5%, 71.6%, and 91.3% for temperate and boreal forests, and 45%, 55%, and 65% for tropical forests. As a sensitivity analysis, we used a threshold of zero to determine reversals. The three major severity scenarios in the main text are ‘low’ severity: *tr45bt385* (a), ‘moderate’ severity: *tr55bt716* (e), and ‘high’ severity: *tr65bt913* (i) (‘tr’: tropical forests, ‘bt’: boreal and temperate forests). Global weighted mean reversal probability by grid area is indicated. Hatched areas indicate the areas with heavily managed forests where probability estimates were interpolated. The inset figures show the kernel density estimates across all forested 8-km grid cells.

**Figure S3.**
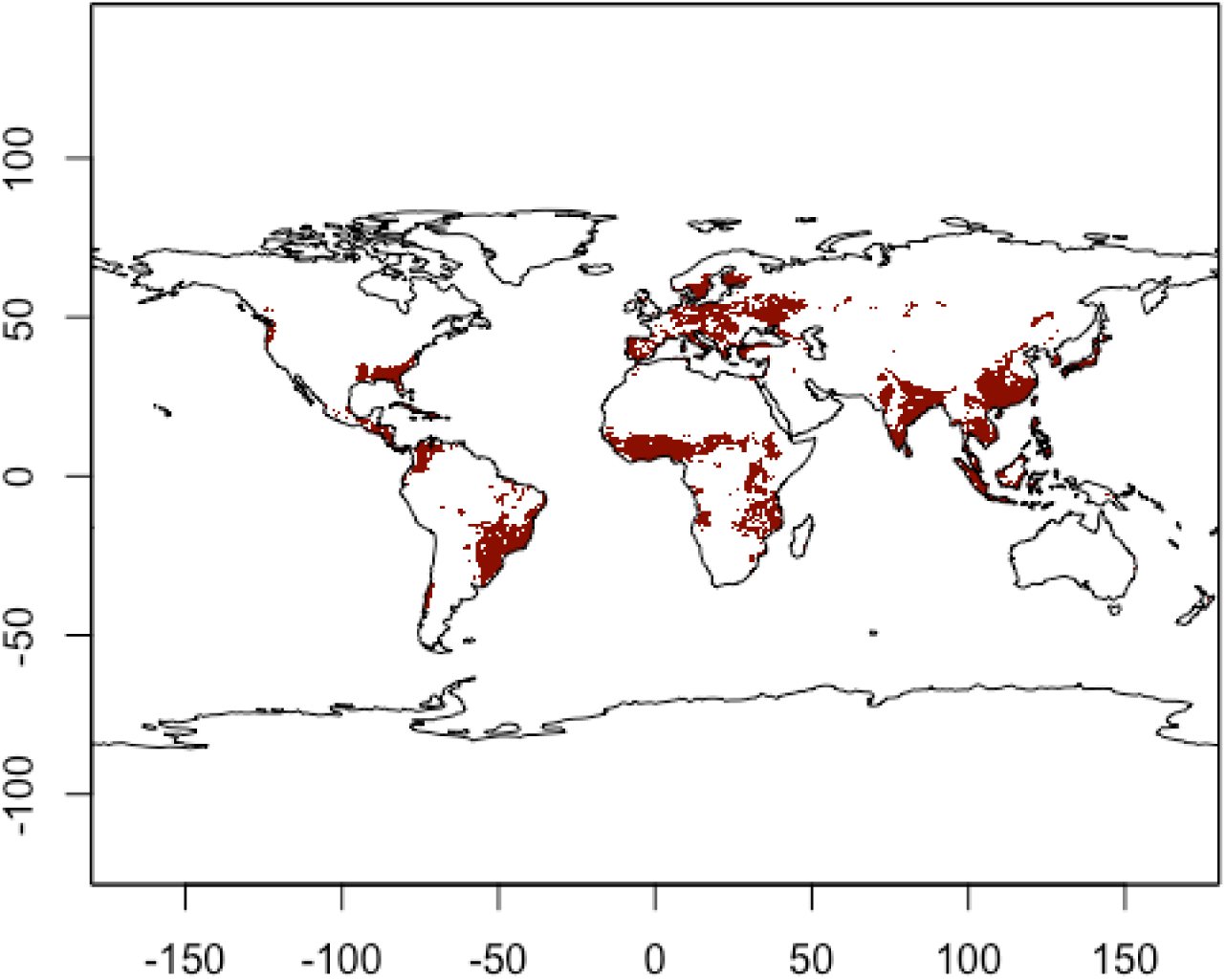
Spatial patterns of the global heavily human-managed forests. The heavily human-managed forests are defined as the forests where the summed fraction of four managed forests (i.e., planted forests (rotation >15 years), plantation forest (rotation ≤15 years), oil palm plantations, and agroforestry) from ref [Lesiv et al., 2022] is > 15%.

**Figure S4.**
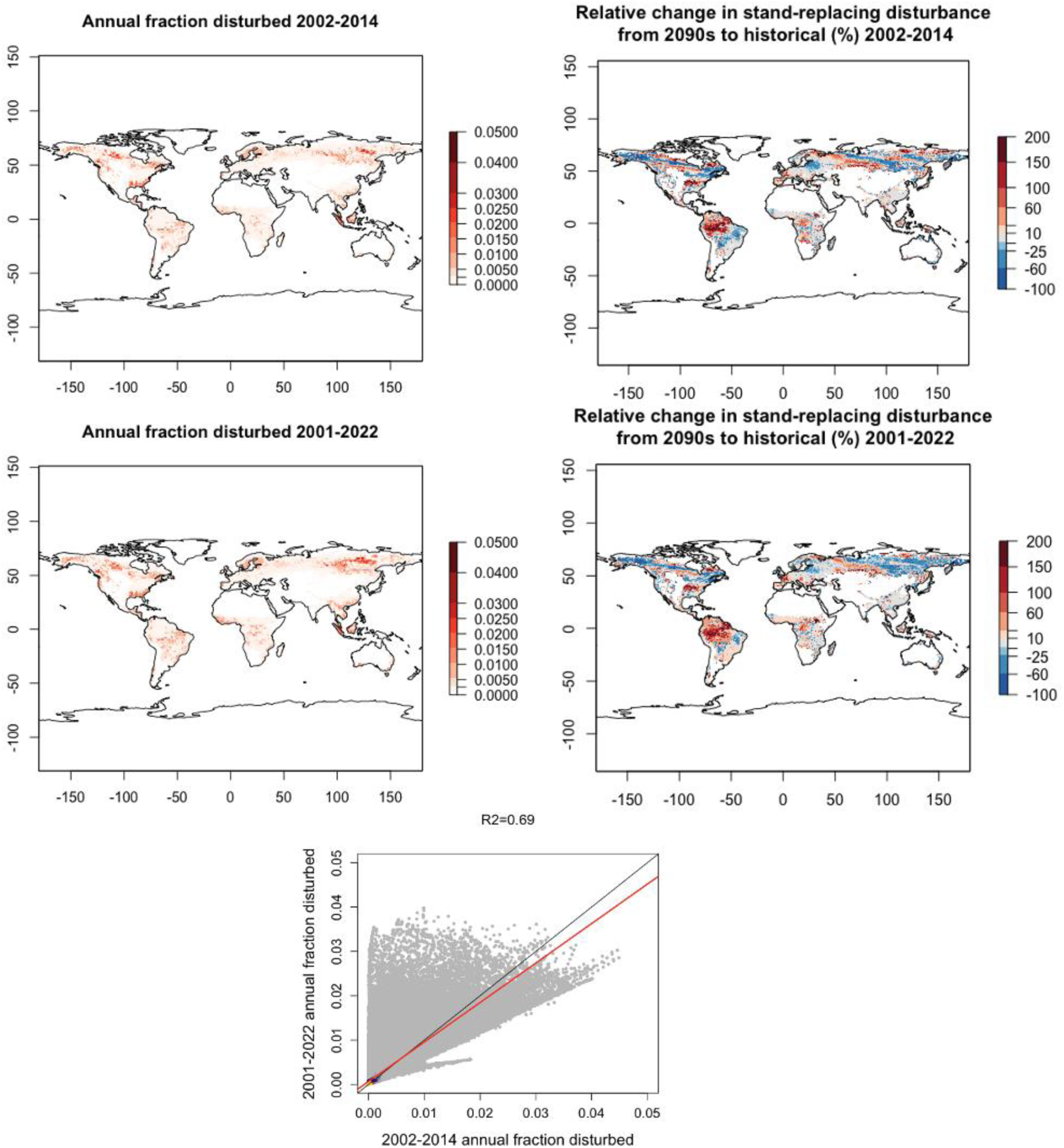
Comparison of the disturbance rate detected from periods of 2001-2022 and 2002-2014. A consistent forest change detection algorithm was employed during the period from 2002 to 2014. Starting in 2015, the change detection algorithm and sensor pool were updated. Here we compared the annual disturbance rate (i.e., fraction disturbed) and relative change in the predicted disturbance rate from the 2090s to the historical between the two disturbance periods detected. The OLS regression result between the two datasets was also shown.

**Figure S5.**
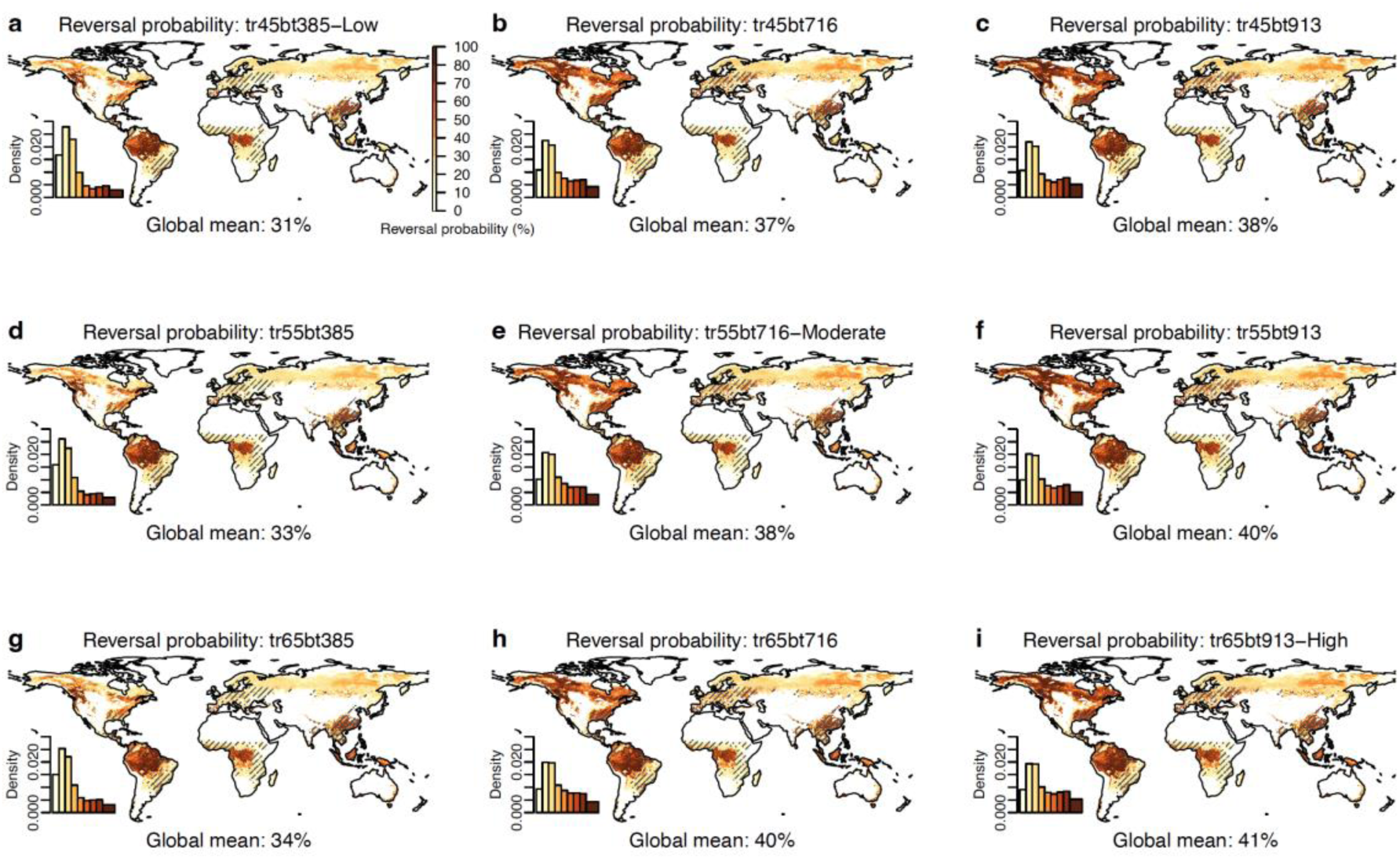
Spatial patterns of the integrated 100-year carbon reversal probability in global forests due to disturbances under the full range of 9 severity scenarios in SSP2-4.5 by combining the conditions where AGB was reduced by 38.5%, 71.6%, and 91.3% for temperate and boreal forests, and 45%, 55%, and 65% for tropical forests. The three major severity scenarios in the main text are ‘low’ severity: *tr45bt385* (a), ‘moderate’ severity: *tr55bt716* (e), and ‘high’ severity: *tr65bt913* (i) (‘tr’: tropical forests, ‘bt’: boreal and temperate forests). Global weighted mean reversal probability by grid area is indicated. Hatched areas indicate the areas with heavily managed forests where probability estimates were interpolated. The inset figures show the kernel density estimates across all forested 8-km grid cells.

**Figure S6.**
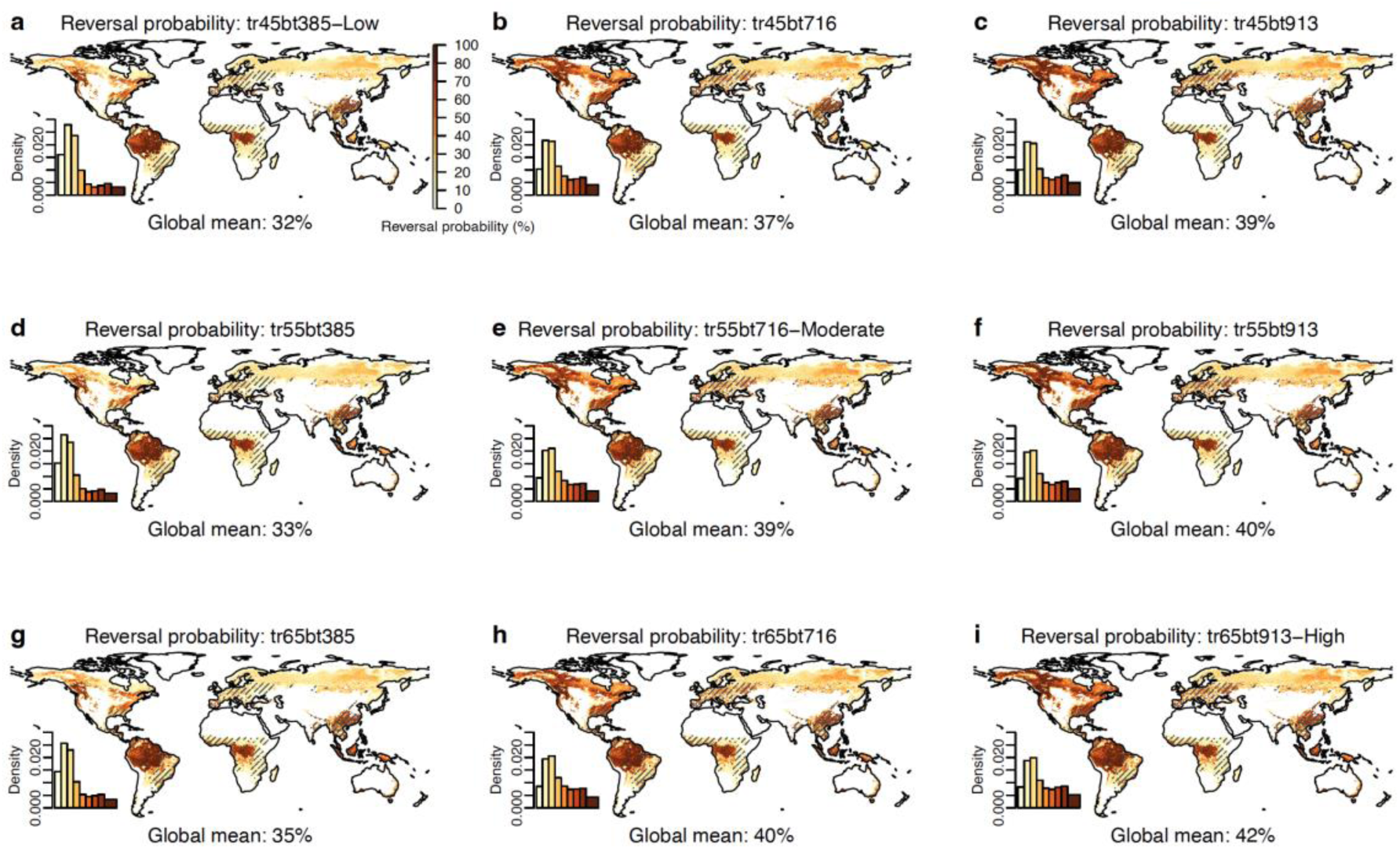
Spatial patterns of the integrated 100-year carbon reversal probability in global forests due to disturbances under the full range of 9 severity scenarios. The same as Figure S4, but under SSP5-8.5.

**Figure S7.**
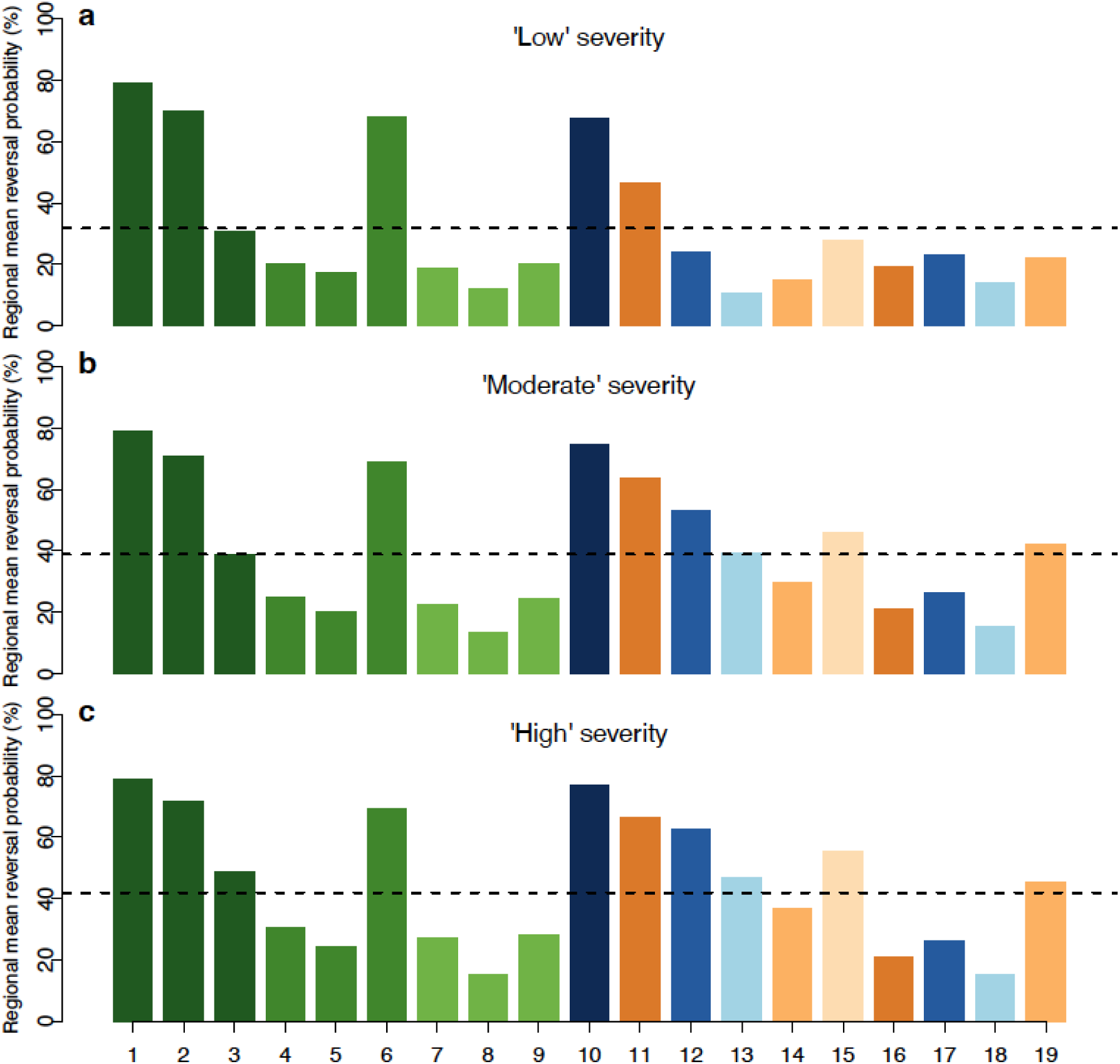
Regional mean probabilities of carbon reversal are provided at a forest ecoregion level under three major severity levels: low (a), moderate (b), and high (c) in SSP5-8.5. The same as Figure 2 in the main text but in a high-emissions scenario.

**Figure S8.**
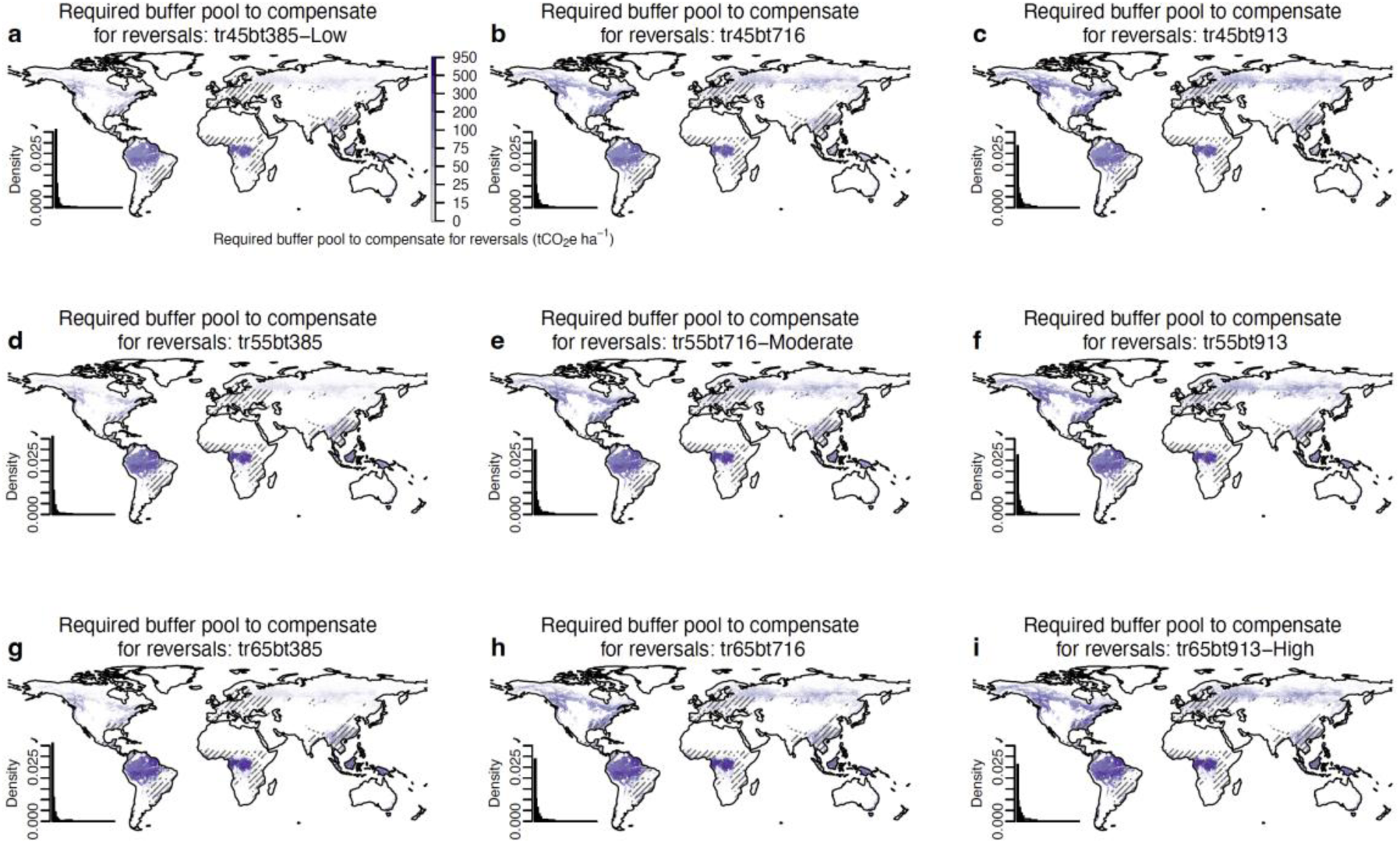
Spatial patterns of the required buffer pool density (tCO_2_e ha^-1^) to compensate for disturbance-driven reversals under the full range of 9 severity scenarios in SSP2-4.5 in global forests by combining the conditions where AGB was reduced by 38.5%, 71.6%, and 91.3% for temperate and boreal forests, and 45%, 55%, and 65% for tropical forests. The three major severity scenarios in the main text are ‘low’ severity: *tr45bt385* (a), ‘moderate’ severity: *tr55bt716* (e), and ‘high’ severity: *tr65bt913* (i) (‘tr’: tropical forests, ‘bt’: boreal and temperate forests). Hatched areas indicate the areas with heavily managed forests where the estimates were interpolated. The inset figures show the kernel density estimates across all forested 8-km grid cells.

**Figure S9.**
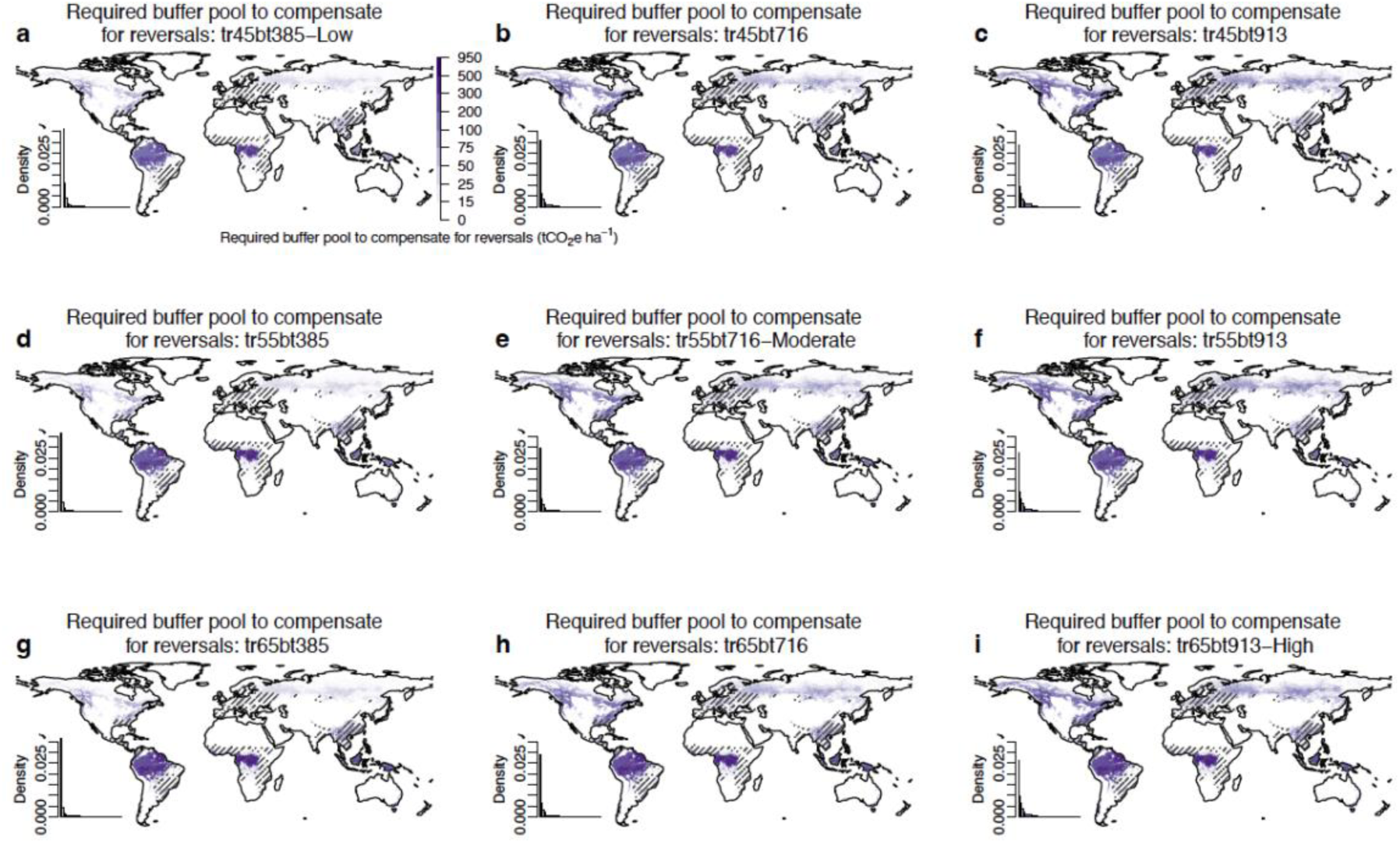
Spatial patterns of the required buffer pool density (tCO_2_e ha^-1^) to compensate for disturbance-driven reversals under the full range of 9 severity scenarios in global forests in SSP5-8.5. The same as Figure S7 but in a high-emissions scenario.

**Figure S10.**
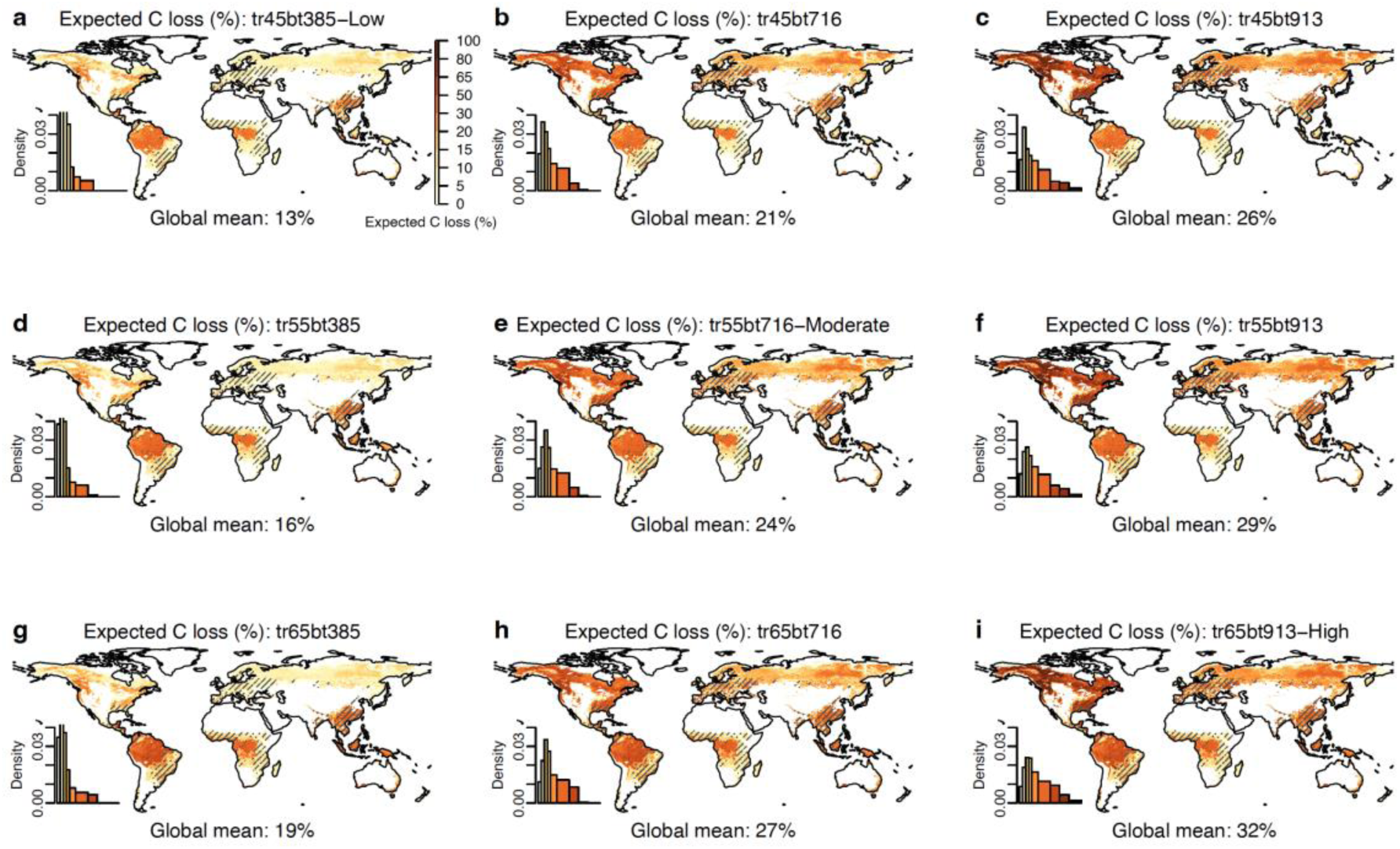
Spatial maps of expected C loss (%) that provides guidance for what buffer pool would be needed to account for disturbances to successfully guarantee total carbon under the full range of 9 severity scenarios in SSP2-4.5 in global forests by combining the conditions where AGB was reduced by 38.5%, 71.6%, and 91.3% for temperate and boreal forests, and 45%, 55%, and 65% for tropical forests. The three major severity scenarios in the main text are ‘low’ severity: *tr45bt385* (a), ‘moderate’ severity: *tr55bt716* (e), and ‘high’ severity: *tr65bt913* (i) (‘tr’: tropical forests, ‘bt’: boreal and temperate forests). Hatched areas indicate the areas with heavily managed forests where the estimates were interpolated. The inset figures show the kernel density estimates across all forested 8-km grid cells. Note that these calculations are made under Accounting Paradigm 2, which makes different assumptions from the results reported in the main text and are not applicable to carbon crediting programs operating under Accounting Paradigm 1.

**Figure S11.**
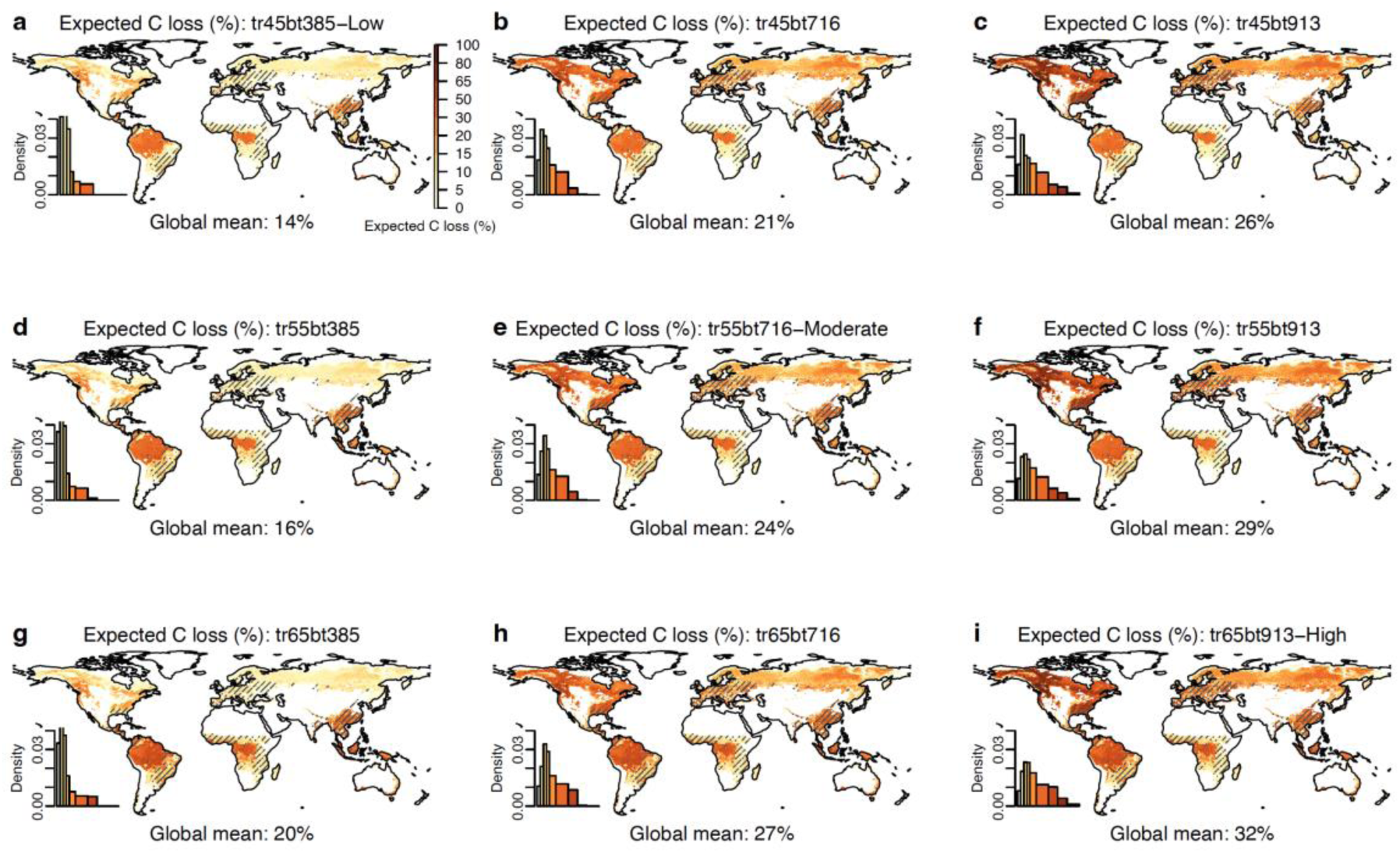
Spatial maps of expected C loss (%) that provides guidance for what buffer pool would be needed to account for disturbances to successfully guarantee total carbon under the full range of 9 severity scenarios in global forests under SSP5-8.5. The same as Figure S9 but in a high-emissions scenario.

**Table S1.** 10 growth functions used to fit the growth curves in global analysis.

| Function No. | Growth function | References |
| --- | --- | --- |
| 1 | $y = a \times \left( \frac{1}{1 + c \times e^{-b \times age} - \frac{1}{c+1}} \right) \times \frac{c+1}{c}$ | [Anderegg et al., 2022; Wu et al., 2023] |
| 2 | $y = a \times age^b$ | [Xu et al., 2023; Yao et al., 2018] |
| 3 | $y = a \times \ln(age + 1)$ | [Zhu et al., 2018] |
| 4 | $y = a \times (1 - e^{-b \times age})^c$ | [Kamo and Yoshimoto, 2013] |
| 5 | $y = a \times e^{\frac{-b}{age}}$ | [Xu et al., 2023; Yao et al., 2018] |
| 6 | $y = \frac{a}{1 + b \times age^{-c}}$ | [Xu et al., 2023; Yao et al., 2018] |
| 7 | $y = \frac{a}{(1 + b \times age^{-d})^c}$ | [Xu et al., 2023; Yao et al., 2018] |
| 8 | $y = \frac{a}{1 + b \times age^{-c}} + d$ | [Kamo and Yoshimoto, 2013] |
| 9 | $y = a \times e^{-b \times age^{-c}}$ | [Kamo and Yoshimoto, 2013] |
| 10 | $y = a \times (1 - e^{-b \times age^c})$ | [Kamo and Yoshimoto, 2013] |

**Table S2.** Fitting performance for the growth curves in 19 ecoregions. The model numnber is the same as Table S1. Asterisk (*) means that the R^2^ is calculated from the best model that is fitted at a broader biome level, otherwise, the R^2^ is calculated from the best model that is fitted at a primary ecoregion level.

| Ecoregion code | Ecoregion name | The best fitted model<sup>#</sup> | R<sup>2</sup> |
| --- | --- | --- | --- |
| 118 | Tundra (Eurasia) | 4 | 0.304120<br>* |
| 117 | Boreal Forest, Shrubland (Eurasia) | 4 | 0.304120<br>* |
| 116 | Temperate Forest, Shrubland (Eurasia) | 4 | 0.304120<br>* |
| 112 | Boreal Forest, Shrubland (North America) | 9 | 0.370668 |
| 113 | Tundra (North America) | 1 | 0.286509 |
| 110 | Conifer Forest (North America) | 4 | 0.465650 |
| 111 | Temperate Forest, Shrubland (North America) | 8 | 0.181471<br>* |
| 119 | Temperate Other Vegetation (Eurasia) | 8 | 0.181471<br>* |
| 114 | Temperate Other Vegetation (North America) | 1 | 0.217365 |
| 106 | Tropical & Subtropical Dry Forest, Shrubland (Asia) | 8 | 0.596683<br>* |
| 101 | Moist Tropical Forest (Americas) | 8 | 0.406996 |
| 109 | Tropical & Subtropical Other Vegetation (Asia) | 9 | 0.227646 |
| 104 | Tropical & Subtropical Dry Forest, Shrubland (Americas) | 3 | 0.212495<br>* |
| 103 | Moist Tropical Forest (Asia) | 2 | 0.369773 |
| 107 | Tropical & Subtropical Other Vegetation (Americas) | 3 | 0.212495<br>* |
| 108 | Tropical & Subtropical Other Vegetation (Africa) | 3 | 0.212495<br>* |
| 105 | Tropical & Subtropical Dry Forest, Shrubland (Africa) | 2 | 0.302940 |
| 102 | Moist Tropical Forest (Africa) | 1 | 0.497176 |
| 115 | Southern Forest (South America) | 5 | 0.333982 |

**Table S3.** CMIP6 models that output mean annual temperature and precipitation, which are further used in the future AGB and disturbance prediction. We list the model name, ensemble member used in this analysis, and reference for each model and data.

| No | Model | Ens. mem. | Reference |
| --- | --- | --- | --- |
| 1 | CESM2 | r4i1p1f1 | [G. Danabasoglu, 2019; Danabasoglu et al., 2020] |
| 2 | CESM2-WACCM | r1i1p1f1 | [Gokhan Danabasoglu, 2019; Danabasoglu et al., 2020] |
| 3 | CMCC-CM2-SR5 | r1i1p1f1 | [Cherchi et al., 2019; Tomas Lovato and Peano, 2020] |
| 4 | CMCC-ESM2 | r1i1p1f1 | [Cherchi et al., 2019; T. Lovato and Peano, 2021] |
| 5 | CNRM-ESM2-1 | r1i1p1f2 | [Seferian, 2019; Séférian et al., 2019] |
| 6 | EC-Earth3-CC | r1i1p1f1 | [Consortium, 2021; Döscher et al., 2021] |
| 7 | EC-Earth3-Veg | r12i1p1f1 | [Döscher et al., 2021; EC-Earth Consortium (EC-Earth), 2019] |
| 8 | EC-Earth3-Veg-LR | r1i1p1f1 | [Döscher et al., 2021; EC-Earth Consortium (EC-Earth), 2020] |
| 9 | MPI-ESM1-2-LR | r1i1p1f1 | [Mauritsen et al., 2019; Wieners et al., 2019] |
| 10 | NorESM2-LM | r1i1p1f1 | [Seland et al., 2019; Seland et al., 2020] |
| 11 | NorESM2-MM | r1i1p1f1 | [Bentsen et al., 2019; Seland et al., 2020] |
| 12 | TaiESM1 | r1i1p1f1 | [W-L Lee and Liang, 2020; W L Lee et al., 2020] |
| 13 | ACCESS-ESM1-5 | r1i1p1f1 | [T. Ziehn et al., 2019; Tilo Ziehn et al., 2020] |
| 14 | BCC-CSM2-MR | r1i1p1f1 | [T Wu et al., 2019; Xin et al., 2019] |
| 15 | CanESM5 | r1i1p1f1 | [N. C. Swart et al., 2019a; N. C. Swart et al., 2019b] |
| 16 | CanESM5-CanOE | r1i1p2f1 | [Neil Cameron Swart et al., 2019; N. C. Swart et al., 2019b] |
| 17 | INM-CM4-8 | r1i1p1f1 | [E. Volodin et al., 2019; Volodin et al., 2017] |
| 18 | INM-CM5-0 | r1i1p1f1 | [*Evgeny Volodin et al., 2019; Volodin et al., 2017*] |
| 19 | IPSL-CM6A-LR | r1i1p1f1 | [*O. Boucher et al., 2019; Olivier Boucher et al., 2020*] |
| 20 | KIOST-ESM | r1i1p1f1 | [*Kim and et al., 2019; Pak et al., 2021*] |
| 21 | MIROC-ES2L | r1i1p1f2 | [*Hajima et al., 2020; Tachiiri et al., 2019*] |
| 22 | UKESM1-0-LL | r1i1p1f2 | [*Good et al., 2019; Sellar et al., 2020*] |

**Table S4.** AGB loss assumptions for each of the three severity scenarios. Note, we also tested six additional scenarios which represented the additional pairwise combinations of different severities for tropical versus extra-tropical forests (e.g., low tropical combined with medium extra-tropical), see Table S5.

| Severity | Tropical<br>(% AGB loss) | Extra-tropical<br>(% AGB loss) |
| --- | --- | --- |
| Low | 45 | 38.5 |
| Moderate | 55 | 71.6 |
| High | 65 | 91.3 |

**Table S5.** The nine disturbance severity scenarios used in global analysis by combining the conditions where AGB was reduced by 38.5%, 71.6%, and 91.3% for temperate and boreal forests, and 45%, 55%, and 65% for tropical forests. The ‘low’, ‘moderate’, and ‘high’ severity scenarios in the main text are indicated and the same as Table S4.

|  |  |  |  |
| --- | --- | --- | --- |
| Boreal & temperate forests (*bt*) | 38.5% AGB lost | 71.6% AGB lost | 91.3% AGB lost |
| Tropical forests (*tr*) |  |  |  |
| 45% AGB lost | ***tr45bt385: low*** | *tr45bt716* | *tr45bt913* |
| 55% AGB lost | *tr55bt385* | ***tr55bt716: moderate*** | *tr55bt913* |
| 65% AGB lost | *tr65bt385* | *tr65bt716* | ***tr65bt913: high*** |

**Table S6.**
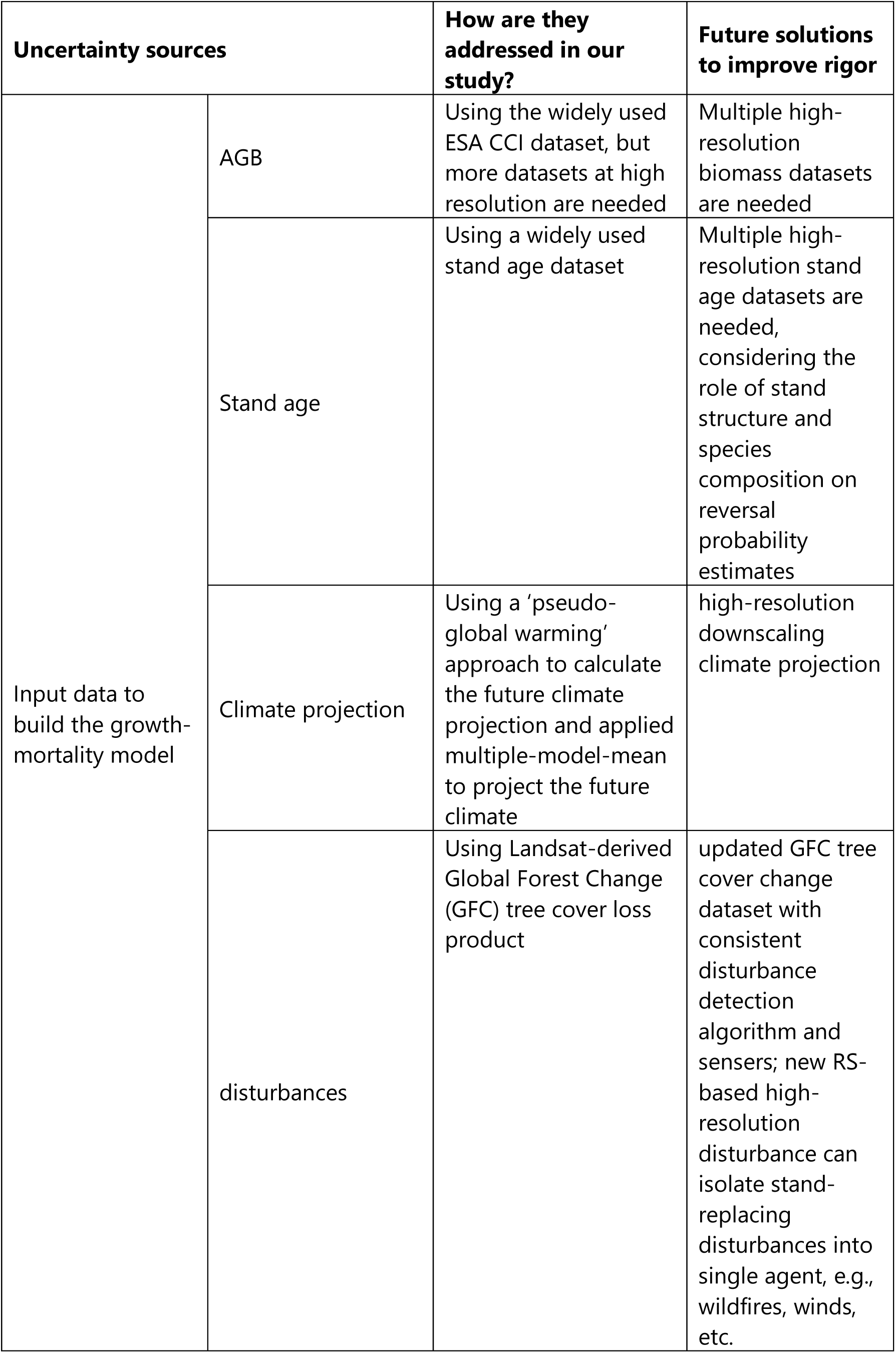
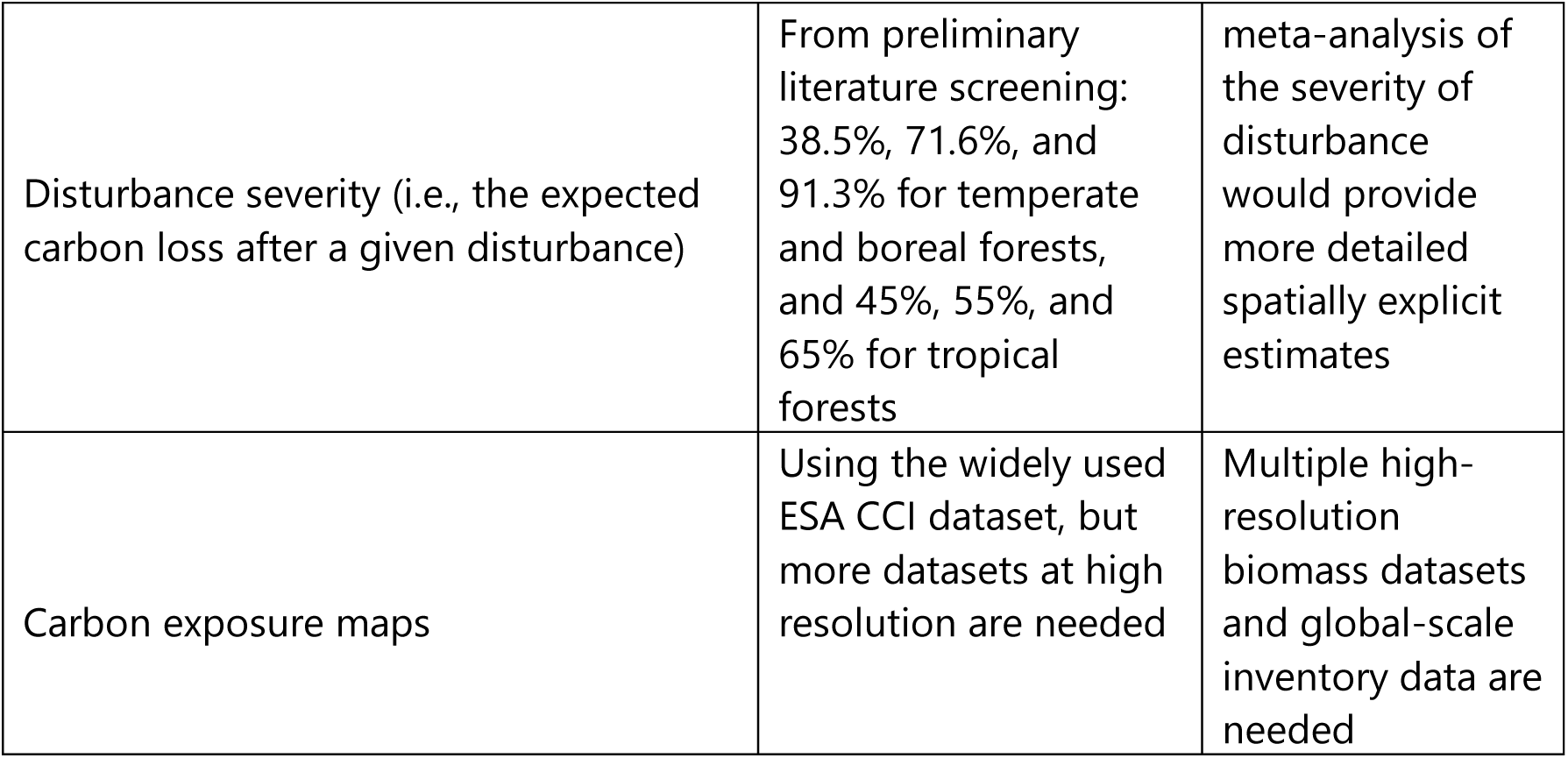
Uncertainties in reversal probability and buffer pool estimates in global analysis and future solutions to improve rigor.

| Uncertainty sources |  | How are they addressed in our study? | Future solutions to improve rigor |
| --- | --- | --- | --- |
| Input data to build the growth-mortality model | AGB | Using the widely used ESA CCI dataset, but more datasets at high resolution are needed | Multiple high-resolution biomass datasets are needed |
|  | Stand age | Using a widely used stand age dataset | Multiple high-resolution stand age datasets are needed, considering the role of stand structure and species composition on reversal probability estimates |
|  | Climate projection | Using a 'pseudo-global warming' approach to calculate the future climate projection and applied multiple-model-mean to project the future climate | high-resolution downscaling climate projection |
|  | disturbances | Using Landsat-derived Global Forest Change (GFC) tree cover loss product | updated GFC tree cover change dataset with consistent disturbance detection algorithm and sensors; new RS-based high-resolution disturbance can isolate stand-replacing disturbances into single agent, e.g., wildfires, winds, etc. |

|  |  |  |
| --- | --- | --- |
| Disturbance severity (i.e., the expected carbon loss after a given disturbance) | From preliminary literature screening: 38.5%, 71.6%, and 91.3% for temperate and boreal forests, and 45%, 55%, and 65% for tropical forests | meta-analysis of the severity of disturbance would provide more detailed spatially explicit estimates |
| Carbon exposure maps | Using the widely used ESA CCI dataset, but more datasets at high resolution are needed | Multiple high-resolution biomass datasets and global-scale inventory data are needed |

